# Extensive population structure highlights an apparent paradox of stasis in the impala (*Aepyceros melampus*)

**DOI:** 10.1101/2024.04.19.590257

**Authors:** Genís Garcia-Erill, Xi Wang, Malthe S. Rasmussen, Liam Quinn, Anubhab Khan, Laura D. Bertola, Cindy G. Santander, Renzo F. Balboa, Joseph O. Ogutu, Patrícia Pečnerová, Kristian Hanghøj, Josiah Kuja, Casia Nursyifa, Charles Masembe, Vincent Muwanika, Faysal Bibi, Ida Moltke, Hans R. Siegismund, Anders Albrechtsen, Rasmus Heller

## Abstract

Impalas are unusual among bovids because they have remained morphologically similar over millions of years – a phenomenon referred to as evolutionary stasis. Here, we sequenced 119 whole genomes from the two extant subspecies of impala, the common (*Aepyceros melampus melampus*) and black-faced (*A. m. petersi*) impala. We investigated the evolutionary forces working within the species to explore how they might be associated with its evolutionary stasis as a taxon. Despite being one of the most abundant bovid species, we found low genetic diversity overall, and a phylogeographic signal of spatial expansion from southern to eastern Africa. Contrary to expectations under a scenario of evolutionary stasis, we found pronounced genetic structure between and within the two subspecies with indications of ancient, but not recent, gene flow. Black-faced impala and eastern African common impala populations had more runs of homozygosity than common impala in southern Africa, and, using a proxy for genetic load, we found that natural selection is working less efficiently in these populations compared to the southern African populations. Together with the fossil record, our results are consistent with a fixed-optimum model of evolutionary stasis, in which impalas in the southern African core of the range are able to stay near their evolutionary fitness optimum as a generalist ecotone species, whereas eastern African impalas may struggle to do so due to the effects of genetic drift and reduced adaptation to the local habitat, leading to recurrent local extinction in eastern Africa and re-colonization from the South.

## Introduction

The impala (*Aepyceros melampus*) belongs to the species-rich Bovidae (Artiodactyla). It has a unique combination of morphological traits that rendered its phylogenetic position challenging until the advent of phylogenomics, which consistently placed it as a basal branch among the antilopine bovids (Antilopinae), together with the genera *Nesotragus* and *Neotragus* (Chen et al., 2019; Hassanin et al., 2012). The impala is the sole extant member of a >10 million-year-old bovid lineage (Chen et al., 2019), and has traditionally been placed in its own tribe, Aepycerotini (Groves & Grubb, 2011). The impala is an example of evolutionary stasis (Eldredge et al., 2005; Vrba, 1984) in a family of otherwise highly radiating taxa, by virtue of its low rates of morphological and taxonomic diversity in the fossil record (Gentry, 1985; Vrba, 1984). The oldest fossils assigned to *Aepyceros* are ≈7 million years old (Geraads, 2019), and specimens assigned to *Aepyceros shungurae*, which appeared in the fossil record between 4 and 3 million years ago, are difficult to distinguish morphologically from modern impalas based on the available cranial material (Gentry, 1985; Vrba, 1984). Such evolutionary stasis is unusual among bovids, making the impala an interesting lineage in which to investigate evolutionary dynamics and to assess possible links between short-term evolutionary dynamics and long-term evolutionary stasis during a time when many other African bovid lineages diversified rapidly.

Evolutionary stasis is characterised as little to no significant morphological changes in a lineage across millions of years (Eldredge et al. 2005). The concept originates from palaeontology, especially in the context of the hypothesis of punctuated equilibrium, which argues that the evolution of most species takes place during rapid bouts of change, which interrupt long periods of stasis (Eldredge & Gould, 1972; Gould & Eldredge, 1977). To our knowledge there are no studies that link population genetic processes with deep-time evolutionary stasis, therefore it is relatively poorly understood which specific combination of population genetic processes characterises evolutionarily stable species, such as the impala. However, theoretical papers have highlighted the roles of stabilizing selection and gene flow in maintaining stasis (Eldredge et al., 2005). First, stabilizing selection tends to maintain populations close to their fitness optimum, a process that is more efficient in larger populations than in smaller ones. If the fitness optimum remains relatively stable, this type of selection – provided it can overcome genetic drift – can lead to evolutionary stasis in the population (S. Estes & Arnold, 2007). Second, even if some local populations deviate from the species-wide fitness optimum or begin adapting to local conditions, gene flow tends to homogenize the species as a whole (Hunt & Rabosky, 2014). Collectively, these factors suggest two testable microevolutionary conditions for evolutionary stasis: (i) stasis is more likely when population structure is limited and populations are not isolated for extended periods, and (ii) stasis is more likely in cases of relatively large effective population sizes (Voje, 2018).

The impala, an ecological generalist, inhabits woodland-grassland ecotones and requires access to water. They exhibit a flexible dietary composition, being able to browse and graze in varying proportions, depending on the season and availability of resources (Fritz & Bourgarel, 2013). Vrba (Vrba, 1980, 1984) posited that the generalist ecology of the impala has been crucial to its morphological stability over extended evolutionary periods. This flexibility enables the impala to cope with variable environments rather than specifically adapt to them. This contrasts with, e.g., the highly variable morphologies found in the bovid specialist tribe Alcelaphini, including species such as the wildebeests and hartebeests. Impalas are usually classified into two subspecies, the common impala (*A. melampus melampus*) and the black-faced impala (*A. melampus petersi*) (Fritz & Bourgarel, 2013). Together, they are distributed throughout the savannah bushlands of southern and eastern Africa, with the common impala having by far the largest distribution and the black-faced impala being restricted to northern Namibia (Figure 1). The species has a combined estimated population size of 2 million individuals, and is the most abundant ungulate in parts of southern Africa, with the black-faced impala amounting to just ≈3300 (perhaps 4300) (Fritz & Bourgarel, 2013). Previous studies using mtDNA concluded that the main genetic structure within the common impala is characterised by a separation of southern from eastern populations, and that eastern populations appeared to have a younger origin (Lorenzen et al., 2006; Nersting & Arctander, 2001). Microsatellite-based studies initially did not find any sign of hybridization between the common and the black-faced impala (Lorenzen & Siegismund, 2004), but a recent study identified several recent hybrids in Etosha National Park and nearby private game reserves (Miller et al., 2020).

**Figure 1:**
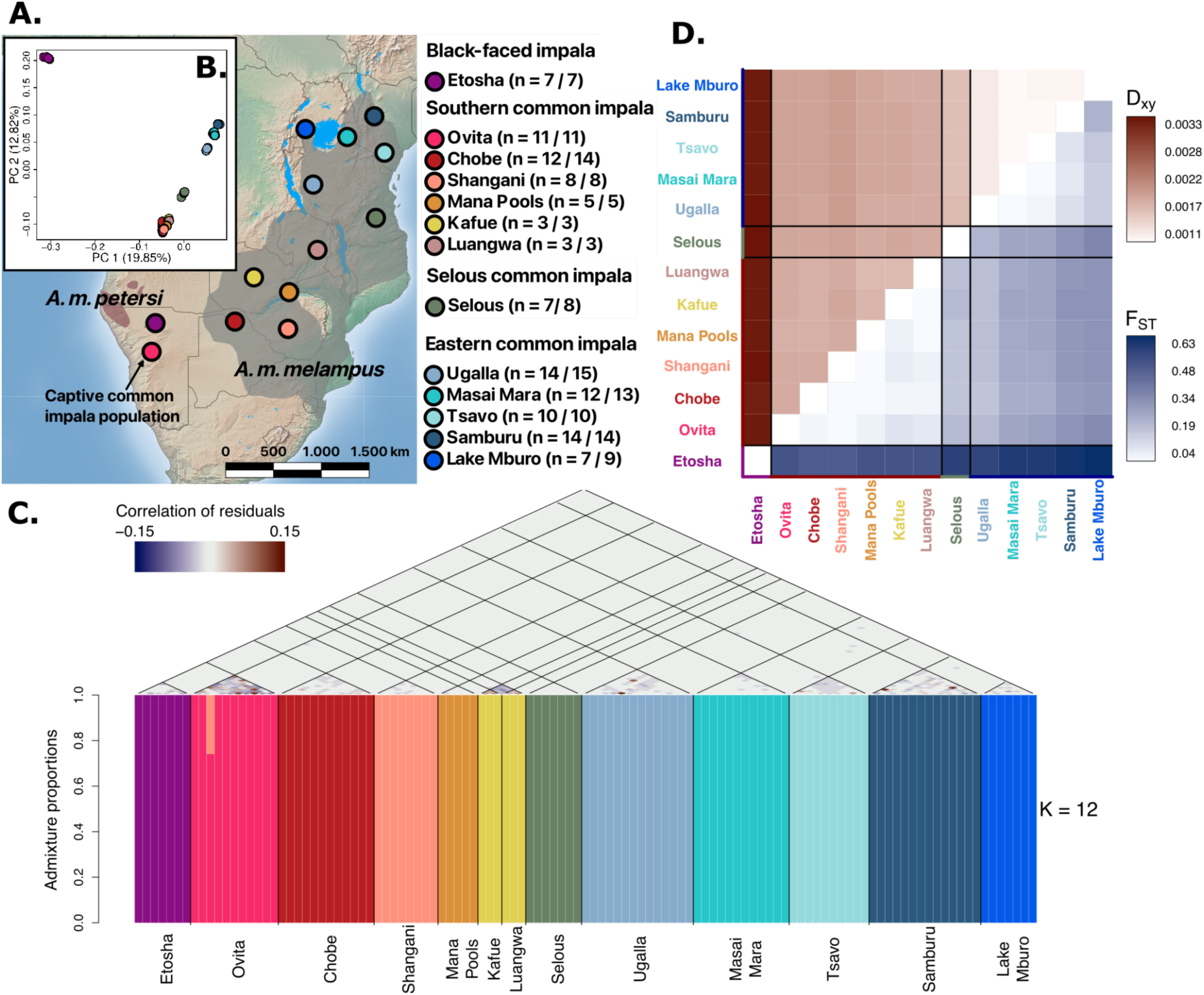
Distribution and population structure of impalas. A. Distribution map and sample localities of impala. Shaded regions indicate the current distribution range of each impala subspecies (IUCN SSC Antelope Specialist Group, 2016). The sample sizes indicated in the legend correspond to the 113 samples kept in the filtered dataset (before the slash) and the initial set of 119 samples sequenced (after the slash), with each locality having at least 1 medium-high depth sample. B. Principal component analysis performed by applying PCAngsd to genotype likelihoods from common SNPs across all impala populations. C. Admixture proportions estimated with NGSadmix, assuming 12 ancestral populations (K=12, lower panel), together with an evaluation of model fit through the correlation of residuals for each pair of individuals estimated with evalAdmix (upper panel). D. Genetic divergence and population differentiation between each population pair, measured respectively with Dxy and with *F*_ST_ estimated using Hudson’s estimator. Both quantities were estimated from the inferred 2-dimensional site frequency spectrum (2DSFS).

To elucidate the evolutionary processes at work within impalas and to understand how these forces might have limited morphological changes from either i) originating and increasing in frequency locally in only one or a few populations, or ii) spreading across the species’ range, we conducted a population genomic study. Specifically, we sequenced 119 whole genomes from both common and black-faced impalas across their range. To address (i), we inferred impala’s genetic structure and admixture, inferred a phylogeographic model to explain the distribution of genetic variation, and considered historical processes such as ancient admixture and changes in population size. For (ii), we estimated historical population sizes and genetic diversity, and compared the efficiency of natural selection in removing new non-synonymous variants that arise in the different populations. We interpret these results in conjunction with fossil record data, exploring various scenarios that could lead to evolutionary stasis, and discuss the implications of our findings for impala conservation.

## Results

We sequenced 119 impala samples from 13 different African localities, comprising 105 samples at low depth (2-4x) and 14 samples at medium depth (7-18x) of genome coverage. Each sampling locality combines low-depth samples and a high-depth sample, which allows the detection of any potential batch effect associated with differences in sequencing depth (Lou & Therkildsen, 2022). Seven of these samples were from black-faced impala, including one medium depth, while the remaining 112 were from common impala (Figure 1, Supplementary Table 1). We mapped the sample reads to both an impala draft reference genome (IMP, (Chen et al., 2019)) and a goat chromosome-level reference genome (ARS1, (Bickhart et al., 2017)), and after strict filtering of sites retained 1.07 and 1.3 Gbp from each respective genome for analyses (Supplementary Table S2). We used preferentially the data mapped to the impala reference genome except for analysis sensitive to reference bias and analyses that require information on chromosomal position of sites, where we used the data mapped to the goat genome because it is an outgroup and it is a chromosome-level genome, respectively (see Method section of each analysis for details). During sample quality control we identified 1 low-depth sample with excessively high sequencing error rates (Supplementary Figure S1) and 5 pairs of duplicate low-depth samples (Supplementary Figure S2). Consequently, we excluded 6 samples from all downstream analyses, resulting in a final dataset of 113 samples (Figure 1A).

### Population structure, recent admixture and genetic diversity

A principal component analysis (PCA) separated the samples from the two subspecies on PC1 and differentiated the southern from the eastern African common impala on PC2 (Figure 1B). Notably, common impalas from Selous were intermediate to southern and eastern Africa on both PC1 and PC2. Higher PCs revealed further population structure within the southern and eastern common impala localities (Supplementary Figure S3). Admixture analyses using NGSadmix (Skotte et al., 2013) corroborated the presence of strong population structure, requiring up to K=12 to obtain an overall good model fit using evalAdmix (Garcia-Erill & Albrechtsen, 2020) (Figure 1C, Supplementary Figures S4 and S5). At K=12, samples from the two Zambian localities, Kafue and Luangwa, showed elevated negative correlation of residuals between them, suggesting they represent separate populations, despite being assigned to the same cluster. Admixture at K > 12 did not converge after 100 independent runs, possibly because the low sample sizes in the two localities prevented the model from assigning each its own cluster at higher K. Finally, the captive common impala population in Ovita showed some noise in individual residual correlations within the population, which could be due to the presence of distantly related individuals and the recent founding of this population, which consists of individuals originally from southern Africa (East, 1998). Otherwise the evalAdmix results are consistent with all sampled localities having distinct and homogeneous genetic ancestries.

Population pairwise *F*_ST_ values confirmed the existence of two broad groups consisting of southern and eastern common impala populations, with Selous showing similar *F*_ST_ (≈0.2) to both groups (Figure 1D). Within the southern populations, *F*_ST_ was relatively low, ranging from 0.04-0.10. In contrast, the eastern group exhibited a slightly wider range of *F*_ST_ values, from 0.04-0.24. This was reversed for D_xy_, where southern populations showed higher D_xy_ (≈0.002) between them than eastern populations (≈0.0011-0.0015). This indicates a more recent divergence, but with greater drift between eastern populations, especially those at the northern extreme of the species’ range, such as in Lake Mburo and Samburu. *F*_ST_ for southern versus eastern populations ranged between 0.18-0.33. Notably, the lower values were mainly found between Selous or Ugalla and the southern populations. Moreover, the D_xy_ between eastern and southern populations were of similar magnitude to the D_xy_ within southern populations, again suggesting a recent divergence. Between black-faced and common impala populations, *F*_ST_ ranged between 0.46-0.61, with the lowest values involving southern populations, while D_xy_ was equally elevated with all common impala populations at ≈0.0033.

Genetic diversity, measured as per-individual heterozygosity, showed a clear geographical trend after removing the low-depth samples with elevated sequencing error rates (Supplementary Figures S1 and S6). The highest genetic diversity values are found in the southernmost populations, Shangani and Chobe, and gradually decline northwards. Surprisingly, despite being geographically restricted and less abundant, the black-faced impala had genetic diversity on par with the southern common impala populations, considerably higher than those in eastern Africa (Figure 2A). This pattern of gradually and monotonically declining genetic diversity in common impala is highly suggestive of a directional expansion (Henn et al., 2019). We explored this further by inferring a population tree with TreeMix. When not allowing for migration, the tree shows a successive split of eastern African common impala populations, roughly coinciding with their geographical distance to southern Africa, with Selous showing the earliest split from the other eastern populations (Figure 2B). This reinforces the notion that the eastern populations likely originated from an expansion of the southern populations.

**Figure 2:**
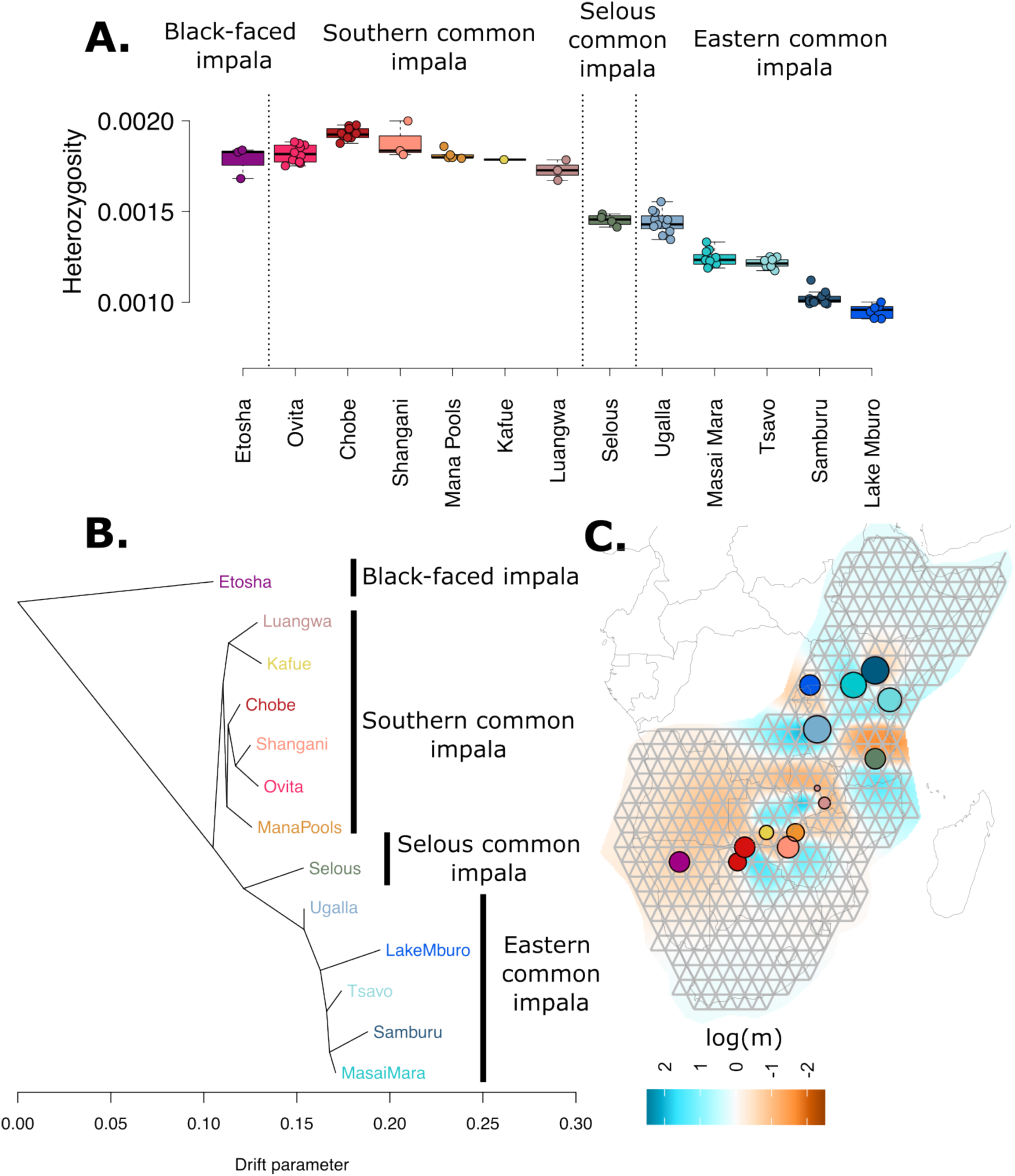
Genetic diversity and phylogeography of impala: A. Per-individual heterozygosity stratified by sampling location for all the impala samples. To minimise bias caused by sequencing errors, we excluded low-depth samples with a sequencing error rate above 0.02% (Supplementary Figure S6). B. Population tree estimated with TreeMix without allowing for any migration events. Labels on the right side show the three main geographical groupings arising from the data. C. Effective migration surface across the species range estimated with EEMS, using data from all the sampled impala populations except the captive common impala population from Ovita Farm. The log(m) values indicate relative migration rates, with lower values (red) corresponding to reduced migration rates and higher values (blue) corresponding to elevated migration rates with respect to the average.

A geography-aware analysis of genetic barriers and connectivity, using EEMS (Estimating Effective Migration Surfaces) on the non-captive populations, confirmed that some parts of the impala range are characterised by barriers to gene flow (Figure 2C). One notable barrier exists between black-faced impala and all other populations. Another separates the southern and eastern common impala populations and a third strong barrier is located north of Selous which possibly geographically overlaps with the Eastern Arc mountain chain. Additional, less apparent barriers identified in the other population structure analyses included one separating Kafue and Luangwa from Mana Pools and Chobe, overlapping with the Zambezi river, and one separating Ugalla from Lake Mburo, overlapping with Lake Victoria. Hence, some barriers correspond with major geographical features that likely serve as physical limits to gene flow, whereas others – notably that between southern and eastern common impala populations, are less clearly linked to present-day geographical features.

### Population divergence and ancestral admixture events

We further used TreeMix to understand ancient gene flow between populations by inferring population trees with admixture between branches. TreeMix has high residuals for up to five admixture events, which suggests abundant gene flow events between populations, including one involving the black-faced impala population from Etosha with its geographically closest southern common impala in Chobe (Supplementary Figure S7). We used D-statistic analyses to further explore the migration events, and used the f-branch statistic implemented in Dsuite (Malinsky et al., 2021) as a visual summary of all combinations of D-statistics with topologies compatible with the TreeMix tree without migration (Figure 2B). This revealed further evidence of multiple admixture events (Figure 3A), including between Selous and Ugalla and the southern populations, and between the two Zambian populations and eastern populations. Moreover, the f-branch statistic also showed evidence of admixture between the common impala in Chobe and the black-faced impala. To further understand this observation, we visualised the D-statistic analyses of the type (((P1,P2),black-faced),Goat), with all possible pairs of common impala populations as P1 and P2. These confirmed that Chobe shares more alleles with the black-faced impala than other common impala populations do, and that southern populations generally share more alleles with black-faced impala than eastern ones (Supplementary Figure S8). Consistent with a Chobe to black-faced impala introgression, a mtDNA tree showed that two black-faced impala individuals carried mtDNA with common impala haplotypes, with Chobe individuals being their closest haplotypes in both cases (Supplementary Figure S9). We used *qpgraph* to quantify the amount of gene flow. To focus on the most important features of the species history and avoid fitting an overly complex model, we chose to include only the following populations: Etosha (black-faced), Chobe (southern common), Shangani (southern common) and Masai Mara (eastern common). We estimated that 16% of black-faced impala ancestry introgressed from Chobe into black-faced impala (Figure 3B). While the admixture graph with Chobe to black-faced introgression fitted the observed f-statistics without any significant deviation, 3 other fundamentally different admixture graphs with a single migration event were equally well supported by the allele sharing data (Supplementary Figure S10). Our choice of the most likely admixture graph out of the 4 candidates was guided by its ability to best explain the presence of common impala mtDNA haplotypes in black-faced impala individuals.

**Figure 3:**
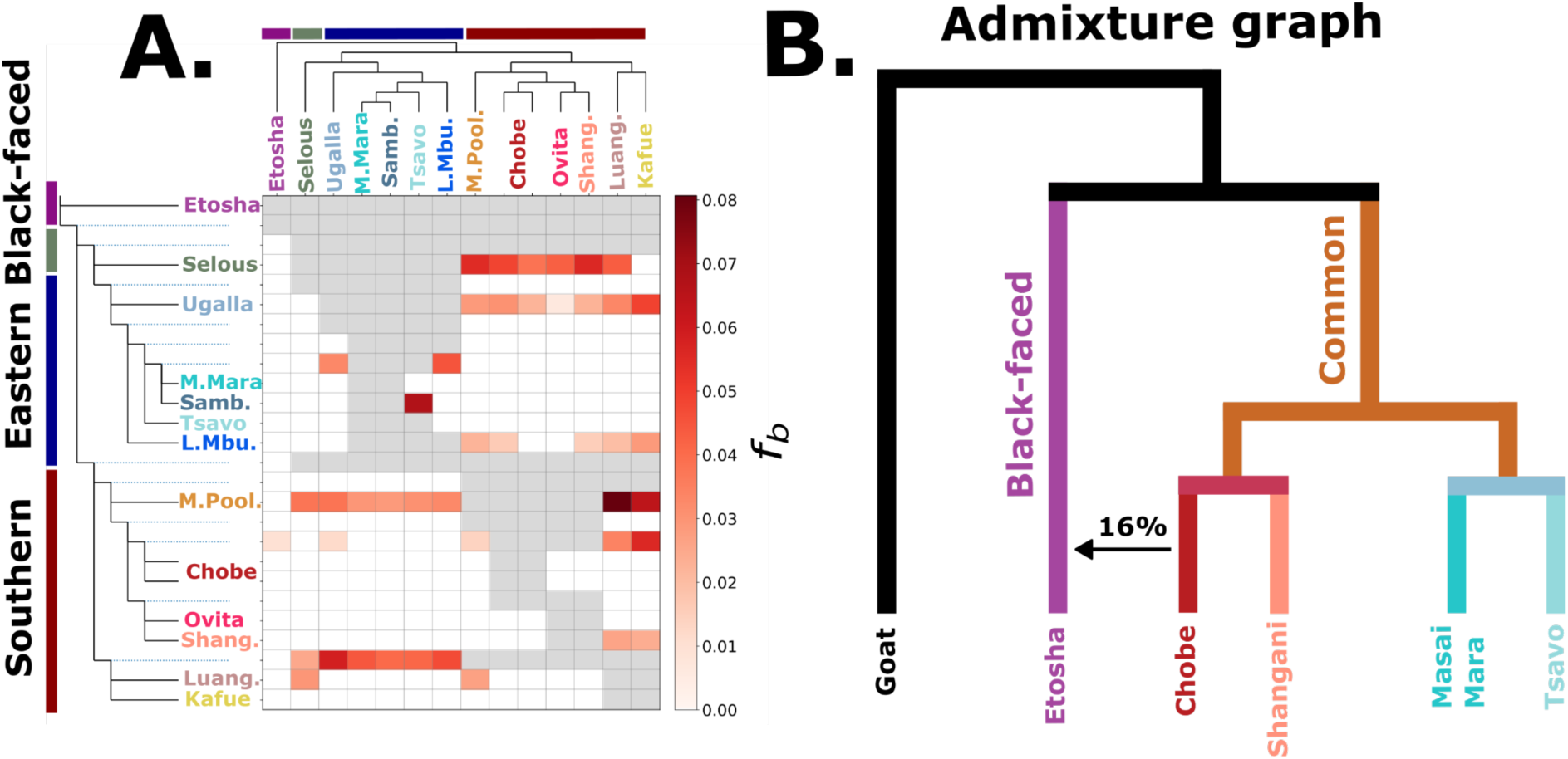
Gene flow between impala populations. A. Summary of all f4 statistics reflecting a significant violation of the population tree without migration (Figure 3B) obtained using the f-branch statistic. Each population is represented by their medium-high depth sample (two for Chobe). B. Admixture graph obtained by applying qpGraph to medium-high depth samples for a subset of the populations. Branches are not drawn to scale. The graph shows a possible scenario of introgression into the black-faced impala from Chobe, its neighbouring common impala population (see Supplementary Figure S10 for other admixture graphs that explain the data equally well).

### Demographic history

We used PSMC to infer the effective population sizes through time from genotype calls of all high and medium-depth samples. To explore the impact of sequencing depth and optimal filtering settings, we also included a previously published high-depth sample from the ruminant genome project (RGP) that was sampled in Laikipia, Kenya (Supplementary Figure S11). PSMC analyses showed a declining trend in population sizes since about 1 million years ago, and highly similar population histories across all impala populations until around 50 kya (Figure 4A). Around 50 kya the southern common impala populations started to increase in size, whereas the eastern as well as the black-faced population continued to decline. We caution that population structure and gene flow can confound population size history inference (Li & Durbin, 2011). Even so, a major split between the demographic histories seems to have occurred around 50 kya. To further corroborate this finding and date the main divergence events in the impala, we carried out demographic history modelling using fastsimcoal2 (Excoffier et al., 2021) with four of the same populations used in the *qpgraph* analysis, excluding the population from Tsavo, Kenya, because it is expected to have a very similar demographic history to the population from Masai Mara and would thus add unnecessary complexity to the model. We tested four different models (Supplementary Figure S12 and Supplementary Tables S2, S4, S5 and S6), and present results for the two with the highest maximum likelihoods. The split time between southern and eastern common impala was estimated at 56 kya, with gene flow occurring at 5 kya, assuming the gene flow was a single admixture pulse (Figure 4B). The divergence time between black-faced and common impala was estimated at 59 kya, with 14% gene flow from Chobe to Etosha occurring around 2 kya. Additionally, we detected highly asymmetric gene flow from Masai Mara to the southern population ancestral to Shangani and Chobe (Figure 4B) and obtained similar results between the southern ancestral population and Masai Mara under continuous gene flow assumptions (Figure 4C). To account for uncertainties in impala’s generation time, we also scaled the demographic history timing with an alternative estimate of generation time. The alternative generation time of 4.03 compared to the would result in younger dating of the different events in impala’s demography, being reduced by a factor of ≈0.71 (Supplementary Figure S13).

**Figure 4:**
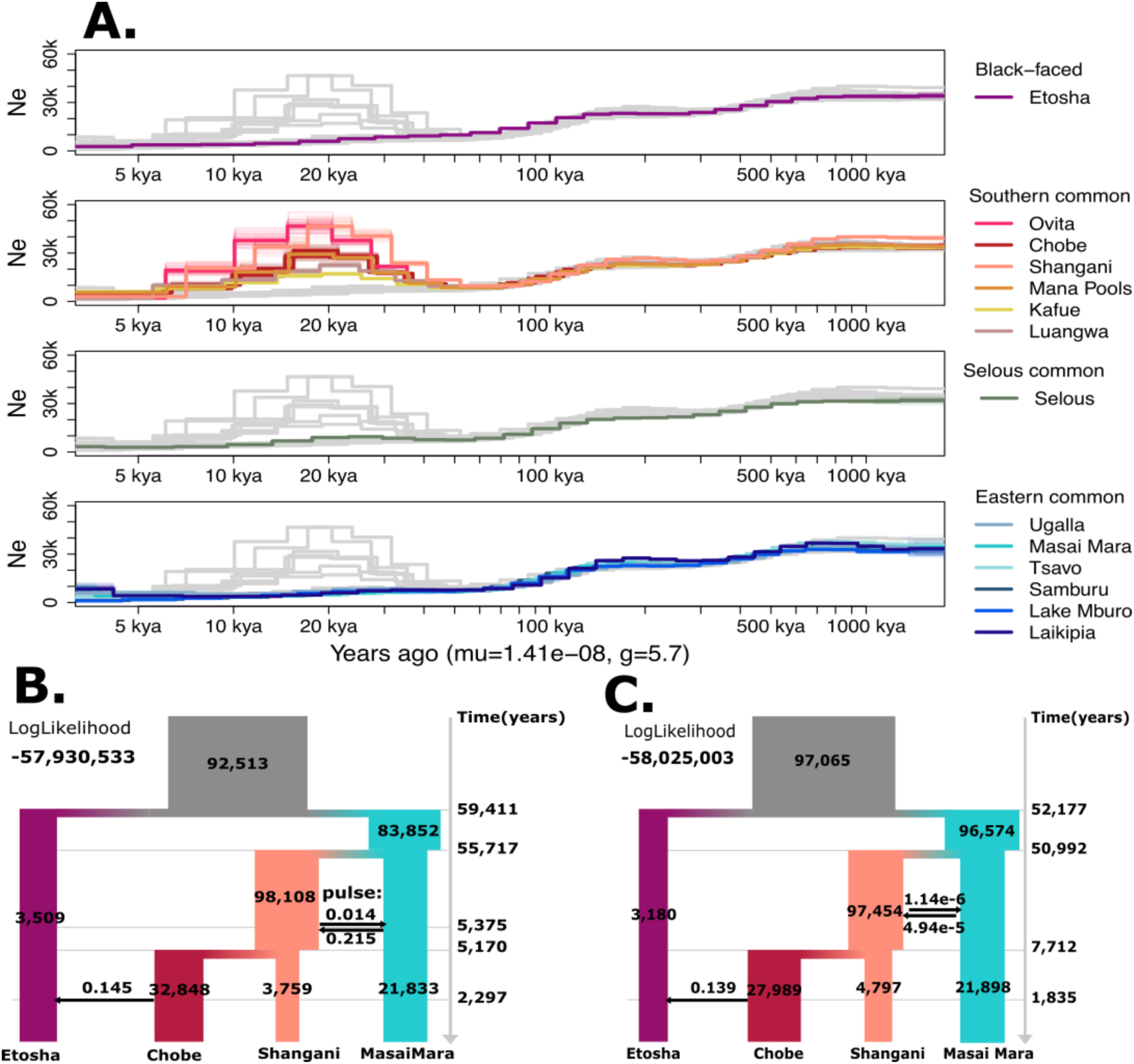
Demographic history of impala: A. Long-term effective population sizes for all impala populations, estimated using the medium-high depth samples and the Pairwise Sequentially Markovian Coalescent (PSMC) algorithm. Each panel highlights coloured samples from each of the three main groups, with bootstrap estimates shown in thinner lines, while the point estimates from non-highlighted samples in each corresponding panel are shown in grey. The high depth sample from the Ruminant Genome Project, collected in Laikipia, Kenya, is included for comparison. B. Joint demographic history for a subset of populations estimated from the 2DSFS with fastsimcoal2, including population split times and admixture events. Time and population sizes are not to scale and the log-likelihood is shown. C. Same model but assuming continuous gene flow between the population ancestral to Shangani and Chobe with Masai Mara.

### Recent population size and strength of selection

To investigate more recent demographic events and their potential effects on genomic diversity, we first inferred the lengths and total proportion of runs of homozygosity (ROHs) in the different individuals. We based these analyses on imputed genotype calls mapped to the goat reference genome, after confirming with a PCA that the imputed data preserved the same main data features as those estimated from genotype likelihoods (Supplementary Figure S14 and Fig 1B). We found that between 1-30% of the genome resides in ROHs across individuals, displaying a clear geographic trend. Specifically, the eastern common impala populations, particularly those in Samburu and Lake Mburo, along with black-faced impala, exhibit progressively more ROHs than the southern common impala populations (Figure 5A). We also found several individuals with a large proportion of their genome in long ROHs (>10 Mbp) in the black-faced impala population from Etosha, the captive common impala population in Ovita and the common impala population in Luangwa, suggesting cases of recent inbreeding within these populations. Linkage disequilibrium (LD) decay curves for each population corroborated this trend, revealing a slower LD decay in the eastern common impala populations, especially in Lake Mburo (Figure 5B), indicative of a reduced effective population size in recent times. The reduced effective population size of the black-faced impala and the eastern common impala can impair the effectiveness of selection. To explore the efficiency of natural selection in removing new functional mutations, which are predominantly harmful, we estimated the ratio of derived alleles at protein coding sites. Specifically, we compared the number of derived alleles at 0-fold degenerate sites, where any substitution alters the encoded amino acid, to those at 4-fold degenerate sites, where substitution preserves the amino acid, across each population. This ratio (Supplementary Figure S15) serves as a proxy for the effectiveness of natural selection in each population (Clemente & Vogl, 2012). This ratio was lower in Selous than in the southern common impala populations, and both were markedly lower than those for the eastern common impala populations and the black-faced impala (Figure 5C). We observed a correlation of the ratio between the total number of derived alleles in the population with the preceding ratio (Supplementary Figure 16), which suggests error rates might bias the ratio by having a higher relative impact on the more evolutionary constrained and less diverse 0-fold degenerate sites. This led us to exclude the population from Shangani, Zimbabwe from the plot, and we warn it might potentially impact the results for the eastern population from Selous, Tanzania, and the black-faced population from Etosha, Namibia.

**Figure 5:**
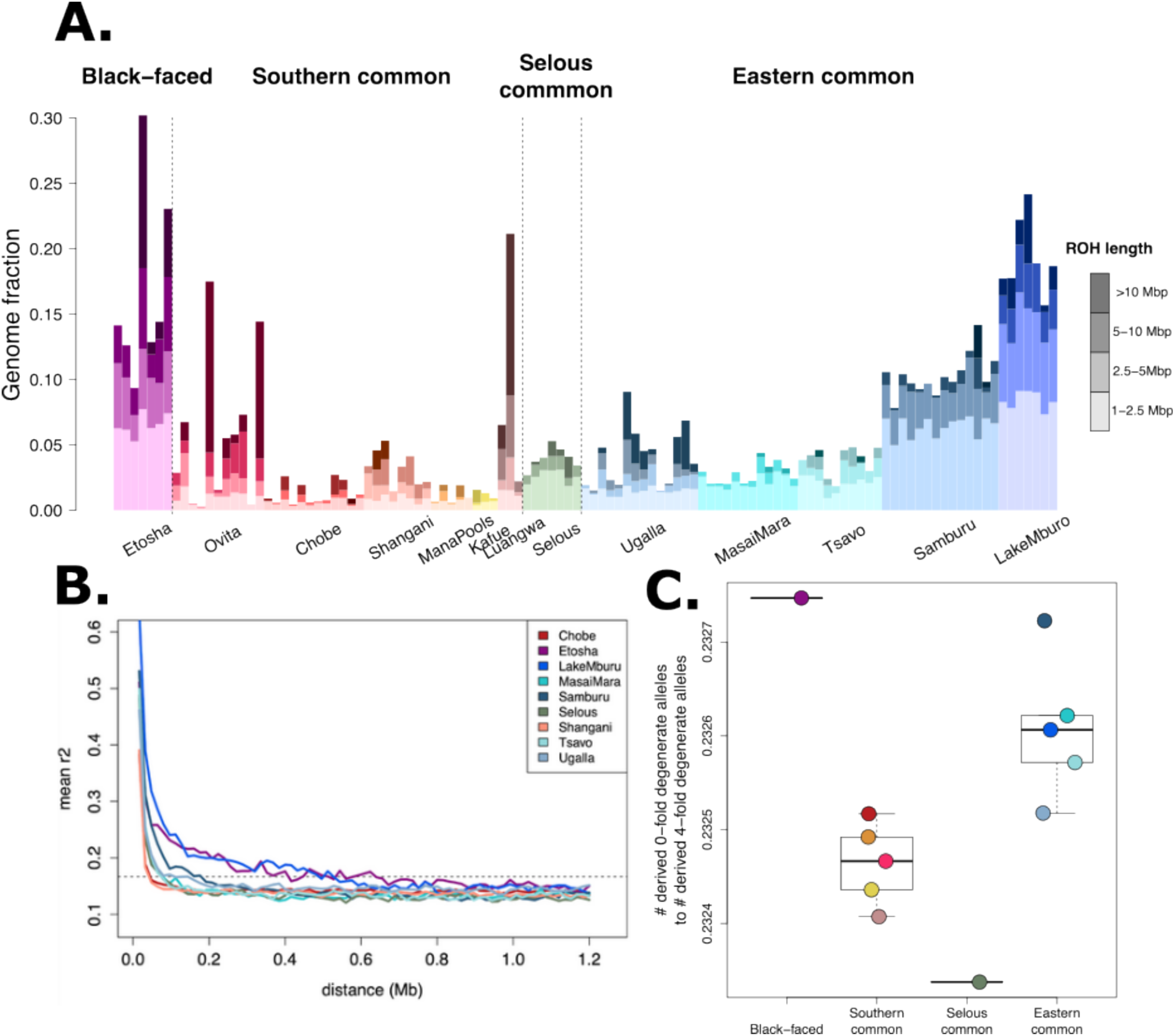
Impact of recent demography in impala genomes. A. Proportion of the genome for each sample covered by runs of homozygosity (ROH). The colour hue distinguishes populations and the colour brightness differentiates the ROH length bins. B. Linkage-disequilibrium decay curves for each population based on N=7 randomly selected individuals from each population. A horizontal dotted guide line is shown at 1/(N-1). C. The ratio of the total number of derived alleles at 0-fold degenerate sites to those at 4-fold degenerate sites within each impala population. The population from Shangani is omitted from this plot due to a likely excess of derived alleles caused by excessive error rates, which could impact the ratio (Supplementary Figure S16).

## Discussion

### A synthesis of the evolutionary processes within the impala

We show that impalas have extensive population genetic structure and can be distinctly classified into three major lineages, each with substantial genetic differentiation: the black-faced impala, the southern common impala and the eastern common impala. Selous, the southernmost population among the eastern common impalas, emerges as a basal branch of this group. It occupies an intermediate position in the PCA plot and has a relatively high F_ST_ (≈0.2) with both the southern and the other eastern populations, indicating it has a unique history and may represent a distinct phylogeographic lineage.

While the estimated divergence between the southern and eastern common impala populations is temporally close to that between black-faced and common impala (≈50-55 kya and ≈52-58 kya, respectively), our models also indicate connectivity post divergence. This suggests a weakly structured population rather than an instantaneous split. The onset of the southern-eastern divergence may have occurred as common impalas began to colonise Eastern Africa from a southern Africa refugium (Lorenzen et al. 2006). This scenario, considering the inherent uncertainties of demographic modelling, accords with the suggestion that common impalas were certainly present in East Africa around 57 kya, and that they may have replaced a morphologically distinct, now-extinct *Aepyceros* population in eastern Africa around this time (Faith et al., 2014). In more recent times, the clear structure and lack of recent admixture between populations imply minimal ongoing gene flow between populations.

The genetic diversity in impalas is exceptionally low (range 0.001-0.002), relative to other African mammals such as the African buffalo (0.0026-0.0051, (Quinn et al., 2023), warthog (0.0025-0.0035, (Garcia-Erill et al., 2022), African leopard (0.0020, (Pečnerová et al., 2021), and waterbuck (0.0030-0.0055, (Wang et al., 2022). This is surprising, given that impalas are both very abundant (East, 1998) and evolutionarily successful, traits typically associated with high population sizes and genetic diversity. A PSMC analysis revealed that impalas have had moderate to low effective population sizes for most of the Pleistocene, which might offer some explanation. One possible explanation is the species’ highly polygynous social structure, characterised by a single territorial male monopolising mating with harems of up to 50-100 females (R. Estes, 1991, 1993). This extreme skew in effective sex ratios could significantly reduce the effective population size. Further research is required to ascertain if this factor alone can explain the surprisingly low genetic diversity in the species. Alternatively, historical confinement to a smaller southern refugium could explain impala’s historically low population sizes.

Despite the evident strong structure and limited recent gene flow, we found signals of extensive ancient admixture across all impala lineages. This includes notable introgression from the common impala to the black-faced impala, with the latter’s ancestry comprising 14-16% common impala genetic ancestry. We inferred this introgression to have occurred around 2 kya, which would explain why it could not be detected using methods aimed at detecting recent admixture on some of the same samples (Lorenzen & Siegismund, 2004). Consistent with this, we found two black-faced individuals carrying common impala mtDNA haplotypes, as was also found in an earlier study (Nersting & Arctander, 2001). We therefore conclude that introgression occurred naturally under climatic conditions that brought the two subspecies into contact, rather than as a result of recent translocations of common impala to Namibia, which has been a concern for the genetic integrity of the subspecies (Lorenzen & Siegismund, 2004; Miller et al., 2020). This introgression might have contributed to the relatively high genetic diversity found in the black-faced impala, despite its recent low population size. However, the Etosha population of the black-faced impala consisted of a small group; approximately 60 individuals, before 1970 (Joubert, 1971). The population was supplemented by 180 individuals from three translocations during 1968-1971 from different areas of its range and three more translocations of a total of 194 from 1970 to 1979 (Green & Rothstein, 1998; Joubert, 1971).

### Are microevolutionary patterns reconcilable with evolutionary stasis in impala?

The pronounced genetic structure and low genetic diversity observed in impalas suggest that they do not meet the two ideal microevolutionary conditions (panmixia and large effective population sizes) for evolutionary stasis. Hence, impalas would seem to exemplify the so-called ‘paradox of stasis’, which describes the discrepancy between the fossil-based inference of morphological stasis and the potential for rapid genetic change observed in many species (Eldredge et al., 2005). In what follows, we integrate our findings on impala population genetics and phylogeography with their fossil record in an attempt to explore and potentially resolve this apparent paradox of stasis in the species.

Evolutionary stasis in impalas has been attributed to impalas being “superbly adapted to their habitat” (Kingdon, 2013). This hypothesis suggests that impalas have reached a fixed-optimum of fitness, where stabilizing selection maintains this peak, and where the fitness optimum remains relatively stable over time, consistent with the notion of prevalent stable adaptive zones suggested by Estes & Arnold (2007). Consequently, most new phenotype-altering mutations – especially those related to these adaptations – would face negative selection. This selection process should be most efficient in large populations and less so in smaller, isolated ones, due to the influence of genetic drift (Hunt & Rabosky, 2014; Voje, 2018). Our results suggest that, indeed, in the eastern common and black-faced impala populations, selection was less efficient in removing functional mutations than in the southern common impala populations. These observations are consistent with the inferred phylogeography indicating that eastern common impala populations descend from a relatively recent spatial expansion from southern Africa, and therefore have experienced more genetic drift. Interestingly, this is also in agreement with morphological variation within the impala (Bastos-Silveira & Lister, 2007), which was subsequently interpreted as evidence that the extant eastern African impala populations originated from a subset of the southern African populations (Reynolds, 2010), and with fossil-based findings suggesting that impalas in eastern Africa were subject to more variable conditions, forcing them to evolve new adaptations or go extinct (Faith et al., 2014; Reynolds, 2007). From this combined evidence, it appears the evolutionary stasis in impalas is consistent with a fixed-optimum hypothesis where southern Africa comprises the core region for impalas. Here, comparatively high effective population sizes (but see above) have allowed populations to stay near a stable fitness peak as an ecotone generalist herbivore under the influence of stabilizing selection. In contrast, eastern African impalas have had more volatile demographic histories, greater morphological variability (Reynolds, 2010) due to less efficient stabilizing selection, and have possibly gone locally extinct at least once (Faith et al., 2014). Our results indicate that, at least during the recent geological past, the extant impala’s optimal habitat was to be found in southern Africa, where it persisted and from where it recolonized eastern Africa, perhaps more than once, after local extirpation events there. Why remains unknown, and climate change, competition with other herbivores, or even hominin-related disturbances are all possible factors worth future investigation. Similarly, we cannot determine for how long southern Africa has acted as a refugium for the impala lineage, nor whether the optimum shifted geographically during the millions of years of its existence. The existence of a Pleistocene southern African refugium for savannah-adapted ungulates with present-day southern and eastern African distributions is supported by phylogeographic evidence from several other species (Lorenzen et al., 2012), including blue wildebeest (Arctander et al., 1999), plains zebra (Pedersen et al., 2018) and common eland (Lorenzen et al., 2010).

Hence, we outline a scenario that could offer a resolution to the paradox of stasis in impalas. The inferred phylogeography, combined with evidence that selection is less efficient in eastern Africa, could explain why impalas manage to stay at a fitness optimum in southern Africa, but fail to do so in eastern Africa. Populations that deviate phenotypically from the fitness optimum may experience higher turnover and local extinction risks (Eldredge et al., 2005), and this may be exacerbated if such “pioneer” populations tend to inhabit areas that are more environmentally volatile. This would cause phenotypically divergent populations to be more temporally transient, and therefore appear infrequently or not at all in the fossil record, leading to an appearance of macroevolutionary stasis despite dynamic microevolutionary changes. Hence, this tentative explanation reconciles a number of key observations in impalas: dynamic microevolutionary change, differential selection across the species range, the existence of a southern African refugium, and the fossil record of the impala. However, this explanation remains a hypothetical reconciliation of the apparent paradox of stasis in impala. It assumes, e.g., that the inferred microevolutionary dynamics – though only informative for the last 100,000 years or so – are representative of evolutionary events on longer time scales, which cannot be taken for granted (Olson, 2024).

### Conservation implications

Our demographic analyses suggest that certain impala populations, in particular the eastern populations and black-faced impala, have maintained small sizes for tens to hundreds of thousands of years. Furthermore, the black-faced population in Etosha and the eastern common impala from Samburu and Lake Mburo show signs of recent population declines, as evidenced by an increase in the proportion of the genome covered by relatively large ROHs (>2 Mb) and in the extent of linkage disequilibrium. In addition, the eastern and black-faced populations have a markedly higher accumulation of functional mutations, indicating less efficient selection. This was especially true for the black-faced population from Etosha, and Samburu common impala, the northernmost population in Africa. Given these findings, it is crucial to monitor these populations at either geographical extreme for potential inbreeding depression and increased genetic load.

The primary conservation concern for the black-faced impala has been the threat of genetic swamping by the much more widespread and abundant common impala. This threat was exacerbated by the translocation of common impala to game farms near Etosha National Park, where the black-faced impala also occurs (Green & Rothstein, 1998). While we found no evidence of recent gene flow between the two subspecies, we identified a relatively large historical introgression event from southern common impala into the black-faced impala, ≈5 kya. This event may have contributed to the black-faced impala’s surprisingly high genetic diversity, despite their historically much lower population sizes than the common impala. Our finding of natural introgression between the two subspecies under certain conditions adds complexity to the task of defining discrete evolutionary units for conservation.

## Materials and methods

### Sample Collection and Laboratory Protocol

We used 119 samples of impala tissue, mainly dried skin, collected during 1994–1999 in Uganda, Kenya, Tanzania, Zambia, Zimbabwe, Botswana and Namibia. All samples, except those from Ovita Game Ranch, were analysed using other genetic markers in previous studies (Lorenzen et al., 2006; Lorenzen & Siegismund, 2004; Nersting & Arctander, 2001) where sample collection and storage is described. The Ovita Ranch population has not previously been analysed, but samples were collected as part of the same project and stored in the same way as the remaining samples.

We used the QIAGEN DNeasy Blood and Tissue Kit (QIAGEN, Valencia, CA, USA) for DNA extraction following the manufacturer’s protocol. We then added RNase to the samples to remove all RNA from the samples. Finally we measured DNA concentrations with Qubit 2.0 Fluorometer (ThermoFisher Scientific) using a dsDNA BR Assay Kit and a Nanodrop One spectrophotometer (ThermoFisher Scientific), and used gel electrophoresis to check the quality of the genomic DNA.

### Sequencing

Library preparation and sequencing was performed at BGI, Shenzhen, China. We used Illumina 150 bp paired-end sequencing. Fourteen samples each representing different sampling locations were sequenced at medium depth (7-18X) using the HiSeq2500 platform while the rest of the samples were sequenced to 2-4X using the NovaSeq platform (Supplementary Table S1).

We also included a high depth sample that was previously published as part of the Ruminant Genome Project (RGP, (Chen et al., 2019)), and that had been used to assemble the impala draft reference genome (BioSample ID SAMN08714480). Based on an initial fastQC screening we selected 5 SRA runs that showed best scores on base composition, quality scores and GC percentage. All runs were short read paired end data sequenced with HiSeq X Ten and with insert sizes of 500 bp (SRR6860888, SRR6860889 and SRR6860891), 600 bp (SRR6860892) and 900 bp (SRR6860896). This data was only used for inference of demographic history with the pairwise sequentially Markovian coalescence (PSMC, see below).

Initial QC was performed on the raw reads using FastQC (Andrews, 2010) and MultiQC (Ewels et al., 2016).

### Mapping

First we used NGSmerge (Gaspar, 2018) on the raw read sequence data to merge all paired reads that overlapped by 11 bp or more. Following this, we separately mapped the merged reads as single-end reads and the non-merged paired-end reads with bwa mem v0.7.17 (Li, 2013) in single-end and paired-end mode, respectively. We then used samtools v. 1.9 to mark and remove duplicate reads, removed low quality alignments (flag -F 3852) and non-merged reads that were not properly mapped (flag -f 3). Finally, we combined both non-merged and merged reads into a single bam file for each sample. We applied this data processing and mapping pipeline using two different reference genomes: the draft impala reference genome IMP (accession GCA_006408695.1, (Chen et al., 2019)) and the goat reference genome ARS1 (refSeq accession GCF_001704415.2, (Bickhart et al., 2017)).

### Common filter settings

For all analyses, we excluded bases with base call quality below 30 and reads with mapping quality below 30. For all analyses that involved estimation of genotype likelihoods with ANGSD (Korneliussen et al., 2014), we used the GATK genotype likelihood model (-gl 2; (McKenna et al., 2010)).

#### Reference genome masks

We masked regions in each of the reference genomes by removing data located in regions that are most likely to contain mapping and genotyping errors (Pečnerová et al., 2021). These were identified with different approaches described below. All approaches were applied with the same settings for both the reference genomes, unless explicitly stated otherwise.

#### Scaffold length

For the impala reference genome, after an initial check of the cumulative amount of sequence contained in scaffolds below a certain length, we excluded all scaffolds shorter than 100 kb, which retained around 92% of all sequences in the reference genome (Supplementary Figure S17). For the goat genome, we discarded all scaffolds not assembled into chromosomes.

#### Repeats

For both reference genomes, we removed all regions annotated as repeats using the corresponding publicly available repeat annotations (Bickhart et al., 2017; Chen et al., 2019).

#### Mappability

We estimated mappability across the genome and excluded all sites with mappability lower than 1. We used genmap (Pockrandt et al., 2020) to infer mappability, using a read length of 100 (-K 100) and a maximum number of mismatches of 2 (-E 2).

#### Sex chromosomes

For most analyses we kept only autosomal chromosomes. For the impala genome, we used a preliminary version of SATC (Nursyifa et al., 2022) to jointly identify sex associated scaffolds and sample sex based on differences in normalised depth between the sexes, and excluded all scaffolds with a significant difference in normalised depth between the two sexes.

#### Depth

We excluded sites with excessively low or high depth. We estimated local depths using ANGSD for two different groupings of samples, one pooling together all low depth samples, excluding only sample 2145 with excessive error rates (Supplementary Figure S1, see methods below), and another pooling together all medium-high depth samples. We then removed from analyses any site with combined depth below 0.5 × median depth or above 1.5 × median depth for either of the two groups.

#### Excess heterozygosity

We estimated a preliminary dataset of genotype likelihoods for the common impala samples after removing samples with high error rates and 5 duplicate samples (see below). We estimated genotype likelihoods with ANGSD, calling variable sites as those with a p-value lower than 1e-6 (-snp_pval 1e-6) and with a minor allele frequency (MAF) higher than 0.05 (--maf 0.05). We used these as input for PCAngsd (Meisner & Albrechtsen, 2018) to conduct a Hardy-Weinberg equilibrium (HWE) test (Meisner & Albrechtsen, 2019), which estimates per site inbreeding coefficients *F* in the range of 1 (total excess of homozygosity) and −1 (total excess of heterozygosity), while using individual frequencies estimated from the first principal components to account for population structure. We used 6 principal components to control for population structure, and only included common impalas because the test cannot account for population structure between highly diverged populations, and thus would not be guaranteed to work well when combining common and black-faced impalas. We identified sites with excess heterozygosity as those showing an F value lower than −0.9 and a p-value for the HWE test lower than 1e-6, and excluded all sites within 10 kbp from them.

### Sample quality control

#### Error rates estimation

We estimated error rates using the ‘perfect sample’ approach implemented in ANGSD (Korneliussen et al., 2014). We used the data mapped to the goat to first generate a consensus fasta sequence for a medium depth common impala sample from Tsavo, Kenya (sample ID 54) and for a medium depth black-faced impala from Etosha, Namibia (sample ID 2076), filtering reads with mapping quality below 30 and bases with base quality below 30. We used each of these two as the ‘error free’ individuals to calculate error rates for all samples, with the goat reference genome as the outgroup. Under the assumption that the perfect individual and the target sample have had equal mutation rates, the method estimates error rates (which are compounded of both sequencing error rates but also errors due to DNA damage) as an excess of mismatches of the sample to the outgroup relative to the mismatches of the perfect sample (Orlando et al., 2013). We initially did the analyses using the goat autosomal chromosomes and no further filtering of sites, using only sample 54 as ‘perfect’. We subsequently repeated the error rate estimation using all site filters and both sample 54 and sample 2076, after observing a systematic reduction in error rates for black-faced impala samples when using the common impala as ‘perfect individual’ (Supplementary Figure S1).

#### Duplicate and relatedness estimation

We used an allele-frequency method (Waples et al., 2019) that estimates three statistics reflecting relatedness between pairs of samples from their 2-dimensional site frequency spectrum (2DSFS), to estimate relatedness between all pairs of samples within each sampling locality. We used ANGSD to estimate site allele frequency (SAF) likelihoods for each individual mapped to the impala reference genome, from which we estimated the 2DSFS between each pair using realSFS (Nielsen et al., 2012).

### Genotype likelihoods

We estimated genotype likelihoods for all 113 impala samples that passed sample filtering, from the data mapped to the impala reference genome after applying all site filters. We called SNPs with ANGSD (Korneliussen et al., 2014) using a SNP p-value of 1e-6 (-snp_pval 1e-6) and a minor allele frequency of 0.05 (-maf 0.05), inferring major and minor alleles from the genotype likelihoods (-doMajorMinor 1).

### Genotype calling

Based on the data mapped to the goat reference genome, we called genotypes for the medium and high depth samples using bcftools 1.13 (Danecek et al., 2021) filtering to minimum base and mapping quality 30. Based on the raw calls, we applied the site filters and removed multiallelic sites as well as all indels. In addition, we masked all genotypes based on less than six reads as well as any heterozygous calls for which either of the two alleles were supported by less than three alternate alleles.

### Imputation

We called genotypes on all 113 samples kept after sample filtering, using imputation to refine the genotype calls. We first used bcftools v1.10 (Danecek et al., 2021) to call genotypes and genotype likelihoods on the data mapped to the goat genome, performing joint calling on all samples with bcftools’ multiallelic caller, keeping only variable sites, and subsequently filtered to remove any site within 10 or less bp of an indel and keep only diallelic SNPs. We then extracted the genotype likelihoods for common SNPs, using ANGSD to estimate allele frequencies and filter sites with maf lower than 0.05, and used the genotype likelihoods as input for imputing genotypes with BEAGLE3 (Browning & Browning, 2009). We subsequently kept only sites with an imputation score (estimated *r*^2^) higher than 0.9, and called the genotype with the maximum posterior probability for each site and individuals, setting to missing sites where the maximum posterior probability was lower than 0.9.

### SFS estimation

We estimated SAF files for all populations using all samples passing sample filtering within each of the populations, using the data mapped to the impala reference genome and using the allele in the reference genome to define the ancestral states, and otherwise using the above described common settings. Using the SAF files, we ran winsfs (Rasmussen et al., 2022) with default parameters to estimate the frequency spectra for all population pairs.

### Inference of population structure

#### Principal component analysis

We used PCAngsd (Meisner & Albrechtsen, 2018) to estimate the covariance matrix between all 113 samples from the common SNP genotype likelihood dataset described above. We used 7 principal components for estimating individual frequencies in PCAngsd’s iterative algorithm to estimate the sample covariance matrix, after testing different numbers of principal components (7, 10 and 12) and visually assessing the principal components obtained. As the eigenvectors of the estimated covariance matrix were not sensitive to variation above 7, we used this value for the analyses presented here.

#### Estimation and evaluation of admixture proportions

We used NGSadmix (Skotte et al., 2013) to estimate admixture proportions from the common SNP genotype likelihood dataset mapped to the impala reference genome, assuming from K=2 to K=13. For each K, we ran independent optimization runs until either the results converged, defined as having the 3 runs with the highest likelihood within 3 or less log likelihood units of each other, or 100 independent runs were reached without convergence, which was the case when assuming K=13. For the results that converged, we evaluated the model fit of the maximum likelihood run for each K by estimating the individual pairwise correlation of residuals with evalAdmix (Garcia-Erill & Albrechtsen, 2020).

### Estimation of D_xy_ and *F*_ST_

Using the population SAF files estimated as previously described, based on the data mapped to the impala genome, we used realSFS to estimate a 2DSFS between each population, that we subsequently used as input to estimate pairwise D_xy_ and *F*_ST_ using Hudson’s estimator (Bhatia et al., 2013) between each population pair with the *sfs* CLI tool (Rasmussen et al., 2024) available at https://github.com/malthesr/sfs.

### Estimation of heterozygosity

We estimated per sample heterozygosity by first estimating individual SAF files with ANGSD from the data mapped to the impala reference genome, with the same settings as previously described. We then used realSFS to estimate the sample SFS, and estimated heterozygosity as the inferred number of heterozygous sites divided by all sites.

### Treemix

Using the SAF files based on the data mapped to the goat reference genome, we called the maximum-likelihood alternative allele count for each site found in every population, leading to approximately 6.8 million input sites for TreeMix (Pickrell & Pritchard, 2012). For each number of migrations between zero and five, we ran TreeMix 25 times with different seeds and chose the best run for each possible migration. In all cases, we specified Etosha as the root.

### EEMS

We used the bed2diffs function, as implemented in Estimating Effective Migration Surfaces (EEMS) (Petkova et al., 2016), to generate an average genetic dissimilarity matrix based on the imputed genotypes of dataset mapped to the goat reference genome, and for each sample set as their coordinates the latitude and longitude of their sampling location (Supplementary Table S1). Then we ran EEMS (Petkova et al., 2016) to assess the decay of genetic similarity in a geospatial context, using 40 million iterations and a burnin of 20 million, and 500 spatial demes spanning East and Southern Africa. We did three independent runs and assessed convergence visually and with the Gelman–Rubin diagnostic in the R package coda (Plummer et al., 2006).

### D-statistics and f-branch

To understand deviations from treeness we summarised it as f-branch statistics using Dsuite (Malinsky et al., 2018, 2021). This was done using the called genotypes from the medium-high depth individuals based on data mapped to the goat reference genome. We use the tree inferred from TreeMix without any migrations.

We also calculated D-statistics with the black-faced impala as P3 and the common impala as P1/P2 using the qpDstat from AdmixTools (Patterson et al., 2012) and summarised the results in a heatmap. We used the called genotypes for the medium-high depth individuals, converted to binary PLINK format using PLINK v1.9 (Chang et al., 2015).

### Admixture graphs

To allow for a more detailed insight into the evolutionary history of impala, ADMIXTOOLS2 (Maier et al., 2023) was used to estimate admixture graphs between populations. f2-scores were first extracted from the PLINK file used to calculate D-statistics (see above) using the ‘extract_f2’ function, using a block length of 4 x 10^6^ cM and retaining only SNPs with a maximal missingness of 0.9 within any given population. The ‘find_graphs’ function was run 500 times to heuristically search for likely graphs describing admixture relationships between populations. In this process, a test score was calculated by optimising a topology from a subset of the f-statistics and comparing all remaining graphs to this topology. Significance testing was then performed using a jackknife approach, only retaining graphs that were non-significantly worse than the graphs with the lowest test score (α=0.05). Graphs that exhibited two admixture events from the same node or a ghost population were subsequently discarded as these could not be distinguished from temporally implausible scenarios. Finally, the number of allowed admixture events in the graph was incremented from zero to the highest number of admixture events where the best test score was not significantly different from the best test score of a less parameterised graph.

### mtDNA analyses

#### mtDNA assembly

We mapped the samples to the impala mitochondrial genome (NCBI accession NC_020675, (Hassanin et al., 2012)), adding also a wildebeest sample (SRA BioSample ID SAMN39917065, (Liu et al., 2024)). All impala and wildebeest samples were mapped to the mitochondrial genome using the same pipeline as for the mapping to the whole genome with bwa mem. We then used ANGSD to produce a consensus fasta sequence for each of the samples, selecting the most common base in each position (-doFasta 2), and removing sites where more than 5% of the reads supported a base different from the most common one.

#### mtDNA tree

The consensus sequences were aligned together with the wildebeest mitogenome sequence used as an outgroup. We used ModelTest-NG v0.1.7 (Darriba et al., 2020) to identify the best-fitting substitution scheme based on the Bayesian Information Criterion. HKY+I+G4 was selected, and with the gamma shape parameter of 0.8698 and the proportion of invariable sites of 0.7043 were used as priors in a phylogenetic tree reconstruction in BEAST v2.7 (Bouckaert et al., 2019). We used a strict molecular clock with a rate parameter of 3.95⨉10^-8^ (Colli et al., 2015) and the Yule tree model. The analysis was run with a MCMC chain length of 10,000,000 samples, logging samples every 1,000 steps and with a pre-burnin of 1,000. We used Tracer v1.7.1 (Rambaut et al., 2018) to verify that the MCMC trace showed convergence and all effective sample sizes had values above 200. TreeAnnotator was used to generate a maximum clade credibility tree based on the common ancestor heights, a 30% burn-in and a 0.7 posterior probability limit.

### Generation time

We used two different generation times, the first is the generation time given by the IUCN (IUCN SSC Antelope Specialist Group, 2016) that was used in the Ruminant Genome Project of (Chen et al., 2019), *T* = 5.7y. However, due to its substantial difference from another published generation time for the impala of 7.3y (Pacifici et al., 2014), we decided to re-estimate the generation time from demographic data. We used the survival rate from (Spinage, 1972). The vector with the survival *l_t_* until year *t* is

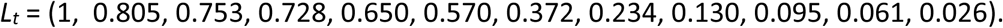

The fecundity vector

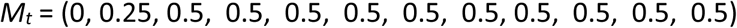

was based on the assumption that the sex ratio in the impala is 1 : 1 and on information obtained from the AnAge Database of Animal Ageing and Longevity (Tacutu et al., 2018) found at https://genomics.senescence.info/species/entry.php?species=Aepyceros_melampus. The generation time was estimated as

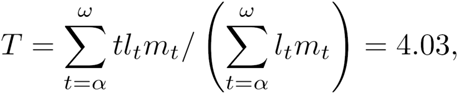

where α is the age of the onset of reproduction and ω is the age of the last reproduction.

### Inference of long-term population size trajectories with PSMC

We used the Pairwise Sequentially Markovian Coalescent (PSMC, (Li & Durbin, 2011)) to estimate population size trajectories through time for the medium-high depth samples. To have a very high depth sample, we used a subset of the short read data from the sample used for the impala genome assembly ((Chen et al., 2019); see Sequencing in Methods section above). We mapped and processed this sample with the sample pipeline as the rest.

Based on the data mapped to the impala genome, we called genotypes separately for each sample with bcftools 1.10, using the consensus caller. We removed sites with a depth lower than either ⅓ of the average sample sequencing depth, or lower than 6 sites if ⅓ of the average was below 6. This minimum depth filter setting was based on an investigation of different settings using subsampled versions of the high depth RGP sample (Supplementary Figure S11B and S11C). We also removed sites with a depth higher than 2 times the average sample depth, and heterozygous sites where there were less than 2 reads supporting either of the two alleles. We then generated psmc input files and ran psmc with default settings. We furthermore performed bootstrapping, by running psmc on bootstrapping mode without splitting, since the scaffolds in the reference genome were already fragmented enough. We scaled the output to years and diploid population sizes assuming a mutation rate of 1.41e-08 per site per generation (Chen et al., 2019) and two different generation times of 5.7 and 4.03 years per generation (see above Generation time section in Methods).

### Inference of joint demographic history with fastsimcoal2

The demographic history of a representative subset of the impala populations (Etosha, Chobe, Shangani and Masai Mara) was further investigated using a coalescent simulation-based method implemented in Fastsimcoal2 v2.7.0.2 (Excoffier et al. 2013). To minimize potential bias arising when determining ancestral allelic states, we used the folded 2dSFS, based on the whole genome 2DSFS estimated with winsfs from the data mapped to the impala reference genome, as previously described. Four plausible demographic models were tested (Supplementary Figure. S12). As the null model, the Model 1 is a scenario without any admixture/migration events and follows a population tree equivalent to the TreeMix tree without any migration edges. The Model 2 scenario adds three admixture events with flexible admixture proportions: one admixture event from Chobe to Etosha and two admixture events happened at the same time before the split between Masai Mara and the ancestor of Chobe and Shangani. The Model 3 models a scenario with one admixture event from Chobe to Etosha and the asymmetric continuous migration between Shangani and Masai Mara. We also considered a Model 4 similar to Model 3, but where asymmetric migration happened continuously before the split between Masai Mara and the ancestor of Chobe and Shangani.

For each model we ran 100 independent Fastsimcoal runs to find the best-fitting parameters yielding the highest likelihood, with 500,000 coalescent simulations per likelihood estimation (-n500000), 100 conditional maximization algorithm cycles (-L100), and minimum 100 observed SFS entry count taken into account in likelihood computation (-C100). A mutation rate of 1.41⨉10^-8^ per site per generation (Chen et al., 2019) and a generation time of 5.7 years (IUCN SSC Antelope Specialist Group, 2016) and 4.03 years (see above section Generation time in Methods) were used to convert model estimates from coalescence units to absolute values (i.e. years).

For the two best models (Model 2 and Model 4), we estimated uncertainty using a non-parametric jackknife approach. We used winsfs to estimate SFS in contiguous blocks of the genome (Rasmussen et al., 2024), producing in total 50 blocks. We then produced 50 leave one out SFS splits by summing the estimated SFS of all except one block each, and estimated again the maximum likelihood parameters for each model, using the maximum likelihood parameters inferred with the full data as initial guesses. For each split we fitted 5 independent runs, and selected the maximum likelihood parameters for each case. Using the 50 maximum likelihood parameters from each split, we estimated standard errors for each parameter using the block jackknife estimator (Busing et al., 1999).

### Runs of homozygosity

We estimated Runs of Homozygosity (ROH) for all samples using the imputed dataset mapped to the goat genome, since the impala reference genome was too fragmented to identify ROHs. We used PLINK 1.9 to call regions in ROH, and after testing different settings used the following: minimum length to call a ROH of 500 kbp (--homozyg-kb 500), minimum SNP density within ROH of 1 SNP per 100 kbp (--homozyg-density 100), used scanning windows of 50 SNPs (--homozyg-window-snp 50), allowing at most 5 heterozygous call within a window in ROH (--homozyg-kb 500), allowing 20 missing SNPs per window in ROH (--homozyg-window-missing 20), and using otherwise default settings. Finally, we manually merged ROHs within 500 kb of each other, after observing a high proportion of ROH within a very short distance most likely due to fragmentation of ROH due to excess missigness or genotyping errors. These settings are quite relaxed, but this was necessary due to a high proportion of spurious heterozygous calls that are unavoidable when mapping to a distant reference genome, which has been shown to often prevent the identification of ROHs (Prasad et al., 2022). Moreover, most of the samples are low depth and imputed without a reference panel, introducing further uncertainty in the genotype calls. However, these limitations affect all the populations equally, and thus, while we advise caution when interpreting the absolute estimated proportion of the genome in ROH for each category, the relative amount of ROHs identified per population can be considered reliable.

### LD-decay

Pairwise LD between sites within populations was estimated using the R package https://github.com/aalbrechtsen/relate (Albrechtsen et al., 2009) and summaries into curves in R as described in https://github.com/aalbrechtsen/LDdecay. We used imputed genotypes mapped to the goat reference genome as input used after removing sites with a minor allele frequency below 5%.

### Efficiency of selection

We used the amount of derived alleles in positions with putatively different functional effects as a proxy for the efficiency of selection. Using the publicly available goat genome annotation, we extracted positions from coding sequences where mutations would change the amino acid sequence of the encoded protein (0-fold degenerate sites), and from those where nucleotide mutations would not alter the amino-acid sequence (4-fold degenerate sites). We assume that mutations in 0-fold degenerate sites would predominantly be neutral or deleterious, while mutations at 4-fold degenerate sites would always be neutral (or would at least be neutral much more often than at 0-fold sites), which is a common assumption made in studies of genetic load and efficiency of selection (Bertorelle et al., 2022). We then estimated population level SAF files from the data mapped to the goat reference genome, separately for each of these two positions using ANGSD with the same settings as previously described, using the allele in the goat genome to assign ancestral state. From the SAF files we estimated the SFS for each population and position type with realSFS. Finally, we used as a proxy of the efficiency of selection within each population, the ratio of the total proportion of derived alleles found within the population at 0-fold degenerate sites to the total proportion of derived alleles found within the population at 4-fold degenerate sites.

## Data availability

All raw sequencing data generated for this study is available in SRA under BioProject accession PRJNA862915. Code used for analyses and visualisations is available at https://github.com/GenisGE/impalascripts.

## Acknowledgements

We thank Amal Al-Chaer for her work on extracting DNA for this study, Peter Arctander for organising the collection of samples, David Moyer for contributing samples to the collection and the African wildlife management authorities for giving permissions and collaborating with Peter Arctander during sample collection. GGE and RH were supported by a Danmarks Frie Forskningsfond Sapere Aude research grant (DFF8049-00098B), RH was further supported by a Carlsberg Young Researcher grant (CF21-0497) and AA, GGE, MSR and KH were supported by Danmarks Frie Forskningsfond (DFF-0135-00211B). We thank the journal editor and two anonymous reviewers for their suggestions that helped improve this manuscript.

## Author contributions

GGE: Conceptualization, Data curation, Formal Analysis, Methodology, Visualization, Writing-original draft, Writing-review & editing; XW: Formal Analysis, Visualization, Writing-review & editing; MSR: Formal Analysis, Methodology, Software, Writing-review & editing; LQ: Conceptualization, Formal Analysis, Writing-review & editing; AK: Conceptualization, Writing-original draft, Writing-review & editing; LDB: Formal Analysis, Visualization, Writing-review & editing; CGG: Formal Analysis, Writing-review & editing; RFB: Formal Analysis, Writing-review & editing; JOO: Conceptualization, Writing-review & editing; PP: Formal Analysis, Writing-review & editing; KH: Methodology, Software, Writing-review & editing; JK: Formal Analysis, Visualization, Writing-review & editing; CN: Formal Analysis, Writing-review & editing; CM: Data curation, Resources, Writing-review & editing; VM: Data curation, Resources, Writing-review & editing; FB: Conceptualization, Writing-review & editing; IM: Conceptualization, Writing-review & editing; HRS: Conceptualization, Data Curation, Resources, Writing-review & editing; AA: Conceptualization, Methodology, Funding acquisition, Project administration, Supervision, Writing-review & editing; RH: Conceptualization, Data Curation, Funding acquisition, Methodology, Project administration, Supervision, Writing-original draft, Writing-review & editing

