## Supplementary tables and figures for "Extensive population structure highlights an apparent paradox of stasis in the impala (*Aepyceros melampus*)"

**Supplementary Table S1**. Sample information table: ID, subspecies, country, locality, coordinate, depth, exclusion criteria

| Sample ID | Subspecies | Country | Population | Latitude | Longitude | Sequencing depth | Exclude Reason |
|---|---|---|---|---|---|---|---|
| 1933 | *A. m. melampus* | Botswana | Chobe | - 18° 42' | 24° 18' | low | |
| 1960 | *A. m. melampus* | Botswana | Chobe | - 18° 42' | 24° 18' | low | |
| 1988 | *A. m. melampus* | Botswana | Chobe | - 18° 42' | 24° 18' | low | |
| 1991 | *A. m. melampus* | Botswana | Chobe | - 18° 42' | 24° 18' | low | Duplicate (with 1988) |
| 2040 | *A. m. melampus* | Botswana | Chobe | - 18° 42' | 24° 18' | low | |
| 2041 | *A. m. melampus* | Botswana | Chobe | - 18° 42' | 24° 18' | medium-high | |
| 2043 | *A. m. melampus* | Botswana | Chobe | - 18° 42' | 24° 18' | low | Duplicate (with 2040) |
| 7825 | *A. m. melampus* | Botswana | Chobe | - 18° 42' | 24° 18' | low | |
| 7830 | *A. m. melampus* | Botswana | Chobe | - 18° 42' | 24° 18' | low | |
| 7695 | *A. m. melampus* | Botswana | Chobe | - 19° 48' | 22° 48' | low | |
| 7829 | *A. m. melampus* | Botswana | Chobe | - 18° 54' | 23° 18' | medium-high | |
| 7826 | *A. m. melampus* | Botswana | Chobe | - 19° 12' | 22° 30' | low | |

| | | | | | | |
|---|---|---|---|---|---|---|
| 7827 | *A. m. melampus* | Botswana | Chobe | - 19° 12' | 22° 30' | low |
| 7828 | *A. m. melampus* | Botswana | Chobe | - 19° 12' | 22° 30' | low |
| 33 | *A. m. melampus* | Kenya | Samburu | 1° 6' | 37° 0' | low |
| 34 | *A. m. melampus* | Kenya | Samburu | 1° 6' | 37° 0' | low |
| 35 | *A. m. melampus* | Kenya | Samburu | 1° 6' | 37° 0' | low |
| 36 | *A. m. melampus* | Kenya | Samburu | 1° 6' | 37° 0' | low |
| 37 | *A. m. melampus* | Kenya | Samburu | 1° 6' | 37° 0' | low |
| 38 | *A. m. melampus* | Kenya | Samburu | 1° 6' | 37° 0' | low |
| 39 | *A. m. melampus* | Kenya | Samburu | 1° 6' | 37° 0' | low |
| 40 | *A. m. melampus* | Kenya | Samburu | 1° 6' | 37° 0' | low |
| 41 | *A. m. melampus* | Kenya | Samburu | 1° 6' | 37° 0' | low |
| 42 | *A. m. melampus* | Kenya | Samburu | 1° 6' | 37° 0' | low |
| 43 | *A. m. melampus* | Kenya | Samburu | 1° 6' | 37° 0' | low |
| 44 | *A. m. melampus* | Kenya | Samburu | 1° 6' | 37° 0' | medium-high |
| 45 | *A. m. melampus* | Kenya | Samburu | 1° 6' | 37° 0' | low |

| | | | | | | |
|---|---|---|---|---|---|---|
| 46 | *A. m. melampus* | Kenya | Samburu | 1° 6' | 37° 0' | low |
| 51 | *A. m. melampus* | Kenya | Tsavo | - 2° 36' | 38° 30' | low |
| 52 | *A. m. melampus* | Kenya | Tsavo | - 2° 36' | 38° 30' | low |
| 53 | *A. m. melampus* | Kenya | Tsavo | - 2° 36' | 38° 30' | low |
| 54 | *A. m. melampus* | Kenya | Tsavo | - 2° 36' | 38° 30' | medium-high |
| 55 | *A. m. melampus* | Kenya | Tsavo | - 2° 36' | 38° 30' | low |
| 56 | *A. m. melampus* | Kenya | Tsavo | - 2° 36' | 38° 30' | low |
| 57 | *A. m. melampus* | Kenya | Tsavo | - 2° 36' | 38° 30' | low |
| 59 | *A. m. melampus* | Kenya | Tsavo | - 2° 36' | 38° 30' | low |
| 60 | *A. m. melampus* | Kenya | Tsavo | - 2° 36' | 38° 30' | low |
| 61 | *A. m. melampus* | Kenya | Tsavo | - 2° 36' | 38° 30' | low |
| 1 | *A. m. melampus* | Kenya | Masai Mara | - 1° 24' | 34° 54' | low |
| 2 | *A. m. melampus* | Kenya | Masai Mara | - 1° 24' | 34° 54' | low |
| 3 | *A. m. melampus* | Kenya | Masai Mara | - 1° 24' | 34° 54' | medium-high |
| 5 | *A. m. melampus* | Kenya | Masai Mara | - 1° 24' | 34° 54' | low |

| | | | | | | | |
|---|---|---|---|---|---|---|---|
| 6 | *A. m. melampus* | Kenya | Masai Mara | - 1° 24' | 34° 54' | low | |
| 7 | *A. m. melampus* | Kenya | Masai Mara | - 1° 24' | 34° 54' | low | |
| 8 | *A. m. melampus* | Kenya | Masai Mara | - 1° 24' | 34° 54' | low | |
| 9 | *A. m. melampus* | Kenya | Masai Mara | - 1° 24' | 34° 54' | low | |
| 10 | *A. m. melampus* | Kenya | Masai Mara | - 1° 24' | 34° 54' | low | |
| 11 | *A. m. melampus* | Kenya | Masai Mara | - 1° 24' | 34° 54' | low | |
| 12 | *A. m. melampus* | Kenya | Masai Mara | - 1° 24' | 34° 54' | low | Duplicate (with 9) |
| 2076 | *A. m. petersi* | Namibia | Etosha | - 18° 48' | 16° 54' | medium-high | |
| 2077 | *A. m. petersi* | Namibia | Etosha | - 18° 48' | 16° 54' | low | |
| 2078 | *A. m. petersi* | Namibia | Etosha | - 18° 48' | 16° 54' | low | |
| 2079 | *A. m. petersi* | Namibia | Etosha | - 18° 48' | 16° 54' | low | |
| 2081 | *A. m. petersi* | Namibia | Etosha | - 18° 48' | 16° 54' | low | |
| 2082 | *A. m. petersi* | Namibia | Etosha | - 18° 48' | 16° 54' | low | |
| 2085 | *A. m. petersi* | Namibia | Etosha | - 18° 48' | 16° 54' | low | |
| 2624 | *A. m. melampus* | Namibia | Ovita | - 21° 36' | 16° 36' | low | |
| 2625 | *A. m. melampus* | Namibia | Ovita | - 21° 36' | 16° 36' | low | |
| 2626 | *A. m. melampus* | Namibia | Ovita | - 21° 36' | 16° 36' | low | |

| | | | | | | | |
|---|---|---|---|---|---|---|---|
| 937 | *A. m. melampus* | Namibia | Ovita | - 21° 36' | 16° 36' | low | |
| 939 | *A. m. melampus* | Namibia | Ovita | - 21° 36' | 16° 36' | low | |
| 942 | *A. m. melampus* | Namibia | Ovita | - 21° 36' | 16° 36' | low | |
| 947 | *A. m. melampus* | Namibia | Ovita | - 21° 36' | 16° 36' | low | |
| 958 | *A. m. melampus* | Namibia | Ovita | - 21° 36' | 16° 36' | low | |
| 959 | *A. m. melampus* | Namibia | Ovita | - 21° 36' | 16° 36' | low | |
| 973 | *A. m. melampus* | Namibia | Ovita | - 21° 36' | 16° 36' | low | |
| 974 | *A. m. melampus* | Namibia | Ovita | - 21° 36' | 16° 36' | low | |
| 3934 | *A. m. melampus* | Tanzania | Selous | - 9° 0' | 37° 24' | medium-high | |
| 8514 | *A. m. melampus* | Tanzania | Selous | - 9° 0' | 37° 24' | low | Duplicate (with 8518) |
| 8515 | *A. m. melampus* | Tanzania | Selous | - 9° 0' | 37° 24' | low | |
| 8516 | *A. m. melampus* | Tanzania | Selous | - 9° 0' | 37° 24' | low | |
| 8517 | *A. m. melampus* | Tanzania | Selous | - 9° 0' | 37° 24' | low | |
| 8518 | *A. m. melampus* | Tanzania | Selous | - 9° 0' | 37° 24' | low | |
| 8519 | *A. m. melampus* | Tanzania | Selous | - 9° 0' | 37° 24' | low | |

| | | | | | | |
|---|---|---|---|---|---|---|
| 8716 | *A. m. melampus* | Kenya | Masai Mara | - 1° 24' | 34° 54' | low |
| 8719 | *A. m. melampus* | Tanzania | Selous | - 9° 0' | 37° 24' | low |
| 4009 | *A. m. melampus* | Kenya | Masai Mara | - 1° 24' | 34° 54' | low |
| 5201 | *A. m. melampus* | Tanzania | Ugalla | - 5° 42' | 32° 0' | low |
| 5454 | *A. m. melampus* | Tanzania | Ugalla | - 5° 42' | 32° 0' | medium-high |
| 5455 | *A. m. melampus* | Tanzania | Ugalla | - 5° 42' | 32° 0' | low |
| 5456 | *A. m. melampus* | Tanzania | Ugalla | - 5° 42' | 32° 0' | low |
| 5457 | *A. m. melampus* | Tanzania | Ugalla | - 5° 42' | 32° 0' | low |
| 5458 | *A. m. melampus* | Tanzania | Ugalla | - 5° 42' | 32° 0' | low |
| 5459 | *A. m. melampus* | Tanzania | Ugalla | - 5° 42' | 32° 0' | low |
| 8485 | *A. m. melampus* | Tanzania | Ugalla | - 5° 42' | 32° 0' | low |
| 8504 | *A. m. melampus* | Tanzania | Ugalla | - 5° 42' | 32° 0' | low |
| 8505 | *A. m. melampus* | Tanzania | Ugalla | - 5° 42' | 32° 0' | low |
| 8708 | *A. m. melampus* | Tanzania | Ugalla | - 5° 42' | 32° 0' | low |
| 8713 | *A. m. melampus* | Tanzania | Ugalla | - 5° 42' | 32° 0' | low |

| | | | | | | | |
|---|---|---|---|---|---|---|---|
| 8715 | *A. m. melampus* | Tanzania | Ugalla | - 5° 42' | 32° 0' | low | |
| 8720 | *A. m. melampus* | Tanzania | Ugalla | - 5° 42' | 32° 0' | low | |
| 2145 | *A. m. melampus* | Uganda | Lake Mburo | - 0° 36' | 31° 0' | low | High error rates |
| 4661 | *A. m. melampus* | Uganda | Lake Mburo | - 0° 36' | 31° 0' | low | |
| 4658 | *A. m. melampus* | Uganda | Lake Mburo | - 0° 36' | 31° 0' | medium-high | |
| 4649 | *A. m. melampus* | Uganda | Lake Mburo | - 0° 36' | 31° 0' | low | |
| 4653 | *A. m. melampus* | Uganda | Lake Mburo | - 0° 36' | 31° 0' | low | Duplicate (with 4649) |
| 4660 | *A. m. melampus* | Uganda | Lake Mburo | - 0° 36' | 31° 0' | low | |
| 4659 | *A. m. melampus* | Uganda | Lake Mburo | - 0° 36' | 31° 0' | low | |
| 4648 | *A. m. melampus* | Uganda | Lake Mburo | - 0° 36' | 31° 0' | low | |
| 4650 | *A. m. melampus* | Uganda | Lake Mburo | - 0° 36' | 31° 0' | low | |
| 4558 | *A. m. melampus* | Zambia | Kafue | - 15° 54' | 25° 54' | medium-high | |
| 2450 | *A. m. melampus* | Zambia | Kafue | - 15° 54' | 25° 54' | low | |
| 2462 | *A. m. melampus* | Zambia | Kafue | - 15° 54' | 25° 54' | low | |
| 2558 | *A. m. melampus* | Zambia | Luangwa | - 12° 6' | 32° 6' | low | |

| | | | | | | |
|---|---|---|---|---|---|---|
| 4555 | *A. m. melampus* | Zambia | Luangwa | - 13° 36' | 31° 48' | medium-high |
| 4556 | *A. m. melampus* | Zambia | Luangwa | - 13° 36' | 31° 48' | low |
| 1617 | *A. m. melampus* | Zimbabwe | Mana Pools | - 15° 54' | 29° 30' | low |
| 1618 | *A. m. melampus* | Zimbabwe | Mana Pools | - 16° 6' | 29° 30' | low |
| 1619 | *A. m. melampus* | Zimbabwe | Mana Pools | - 16° 6' | 29° 30' | low |
| 1620 | *A. m. melampus* | Zimbabwe | Mana Pools | - 16° 6' | 29° 30' | low |
| 1621 | *A. m. melampus* | Zimbabwe | Mana Pools | - 16° 6' | 29° 30' | medium-high |
| 1508 | *A. m. melampus* | Zimbabwe | Shangani | - 18° 48' | 28° 18' | medium-high |
| 1509 | *A. m. melampus* | Zimbabwe | Shangani | - 18° 48' | 28° 18' | low |
| 1510 | *A. m. melampus* | Zimbabwe | Shangani | - 18° 48' | 28° 18' | low |
| 1511 | *A. m. melampus* | Zimbabwe | Shangani | - 18° 48' | 28° 18' | low |
| 1514 | *A. m. melampus* | Zimbabwe | Shangani | - 18° 48' | 28° 18' | low |
| 1515 | *A. m. melampus* | Zimbabwe | Shangani | - 18° 48' | 28° 18' | low |
| 1516 | *A. m. melampus* | Zimbabwe | Shangani | - 18° 48' | 28° 18' | low |
| 1517 | *A. m. melampus* | Zimbabwe | Shangani | - 18° 48' | 28° 18' | low |

**Supplementary Table S2**. Site filters summary. Proportion of the genome removed upon genome filtering based on different criteria, for each of the two reference genomes used.

| | Impala draft reference genome | | Goat genome reference | |
|---|---|---|---|---|
| | Total bp removed | Proportion removed | Total bp removed | Proportion remove |
| RepeatMasker | 900,788,476 | 0.39 | 956,483,991 | 0.39 |
| Mappability | 219,583,124 | 0.09 | 187,595,633 | 0.08 |
| Excess heterozygosity | 94,372,549 | 0.04 | 329,059,218 | 0.13 |
| Depth | 194,098,696 | 0.08 | 570,974,263 | 0.23 |
| All filters combined | 1,070,741,734 | 0.46 | 1,302,327,747 | 0.53 |

**Supplementary Table S3.** Inferred parameters of demographic history between Impala populations for Model 1 (null model): g=5.7 years, mutation rate=1.41e-8, see Supplementary Figure S12 for model visualisation. The log likelihood under the shown model parameters is **-58,748,267.**

| **Parameters** | | | | **Point estimation** |
|---|---|---|---|---|
| **Effective population Sizes (number of diploid individuals)** | **Ancestral** | Chobe&Shangani | NANC1 | 97380 |
| | | Shangani&MasaiMara | NANC2 | 93923 |
| | | Etosha&MasaiMara | NANC3 | 97372 |
| | **Population** | Etosha | NPOP0 | 6340 |
| | | Chobe | NPOP1 | 6053 |
| | | Shangani | NPOP2 | 1883 |
| | | MasaiMara | NPOP3 | 17946 |
| **Time (years)** | **Divergence event** | Chobe&Shangani | TDIV1 | 3232 |
| | | Shangani&MasaiMara | TDIV2 | 40276 |
| | | Etosha&MasaiMara | TDIV3 | 48256 |

**Supplementary Table S4.** Inferred parameters of demographic history between Impala populations for best model - Model 2 (single pulse admixture event before split): g=5.7 years, mutation rate=1.41e-8, see Supplementary Figure S12 model visualisation. The log likelihood under the shown model parameters is **-57,930,533.**

| **Parameters** | | | | **Point estimation** | **Standard error** |
|---|---|---|---|---|---|
| **Effective population Sizes (number of diploid individuals)** | **Ancestral** | Chobe&Shangani | NANC1 | 98108 | 38824 |
| | | Shangani&MasaiMara | NANC2 | 83852 | 76130 |
| | | Etosha&MasaiMara | NANC3 | 92513 | 20378 |
| | **Population** | Etosha | NPOP0 | 3509 | 576 |
| | | Chobe | NPOP1 | 32848 | 36333 |
| | | Shangani | NPOP2 | 3759 | 6260 |
| | | MasaiMara | NPOP3 | 21833 | 9068 |
| **Time (years)** | **Admixture event** | Chobe&Etosha | TADM1 | 2297 | 2331 |
| | | Shangani&MasaiMara | TADM2 | 5375 | 6275 |
| | **Divergence event** | Chobe&Shangani | TDIV1 | 5170 | 6717 |
| | | Shangani&MasaiMara | TDIV2 | 55717 | 7639 |
| | | Etosha& MasaiMara | TDIV3 | 59411 | 8640 |
| **Admixture Parameters** | | Chobe to Etosha | PADM1 | 0.145 | 0.012 |
| **Admixture proportions** | | MasaiMara to Shangani | PADM2 | 0.215 | 0.031 |
| | | Shangani to MasaiMara | PADM3 | 0.014 | 0.003 |

**Supplementary Table S6.** Inferred parameters of demographic history between Impala populations for Model3 (continuous migration before split): g=5.7 years, mutation rate=1.41e-8, see Supplementary Figure 12 for model visualisation. The log likelihood under the shown model parameters is **-58,128,757.**

| **Parameters** | | | | **Point estimation** |
|---|---|---|---|---|
| **Effective population Sizes (number of diploid individuals)** | **Ancestral** | Chobe & Shangani | NANC1 | 116036 |
| | | Shangani&MasaiMara | NANC2 | 72070 |
| | | Etosha&MasaiMara | NANC3 | 55774 |
| | **Population** | Etosha | NPOP0 | 397 |
| | | Chobe | NPOP1 | 160063 |
| | | Shangani | NPOP2 | 16038 |
| | | MasaiMara | NPOP3 | 21565 |
| **Time (years)** | **Admixture event** | Chobe&Etosha | TADM1 | 661 |
| | **Divergence event** | Chobe&Shangani | TDIV1 | 39917 |
| | | Shangani&MasaiMara | TDIV2 | 57832 |
| | | Etosha&MasaiMara | TDIV3 | 58774 |
| **Admixture Parameters** | | Chobe&Etosha | PADM1 | 0.296 |
| **Migration Rates** | | MasaiMara to Shangani | mSM | 3.33e-4 |
| | | Shangani to MasaiMara | mMS | 3.85e-4 |

**Supplementary Table S6.** Inferred parameters of demographic history between Impala pops for Model4 (migration happened before split): g=5.7 years, mutation rate=1.41e-8, see Supplementary Figure 12 for model visualisation. The log likelihood under the shown model parameters is **-58,025,003.**

| **Parameters** | | | | **Point estimate** | **Standard error** |
|---|---|---|---|---|---|
| **Effective population Sizes (number of diploid individuals)** | **Ancestral** | Chobe&Shangani | NANC1 | 97454 | 4445 |
| | | Shangani&MasaiMara | NANC2 | 96574 | 13203 |
| | | Etosha&MasaiMara | NANC3 | 97065 | 11252 |
| | **Population** | Etosha | NPOP0 | 3180 | 355 |
| | | Chobe | NPOP1 | 27989 | 12943 |
| | | Shangani | NPOP2 | 4797 | 4035 |
| | | MasaiMara | NPOP3 | 21898 | 15405 |
| **Time (years)** | **Admixture event** | Chobe&Etosha | TADM1 | 1834 | 2900 |
| | **Divergence event** | Chobe&Shangani | TDIV1 | 7712 | 7744 |
| | | Shangani&MasaiMara | TDIV2 | 50992 | 20386 |
| | | Etosha&MasaiMara | TDIV3 | 52178 | 19080 |
| **Admixture Parameters** | | Chobe&Etosha | PADM1 | 0.139 | 0.084 |
| **Migration Rates** | | MasaiMara to Shangan&Chobe | mSM | 4.942e-5 | 3.137e-6 |
| | | Shangani&Chobe to MasaiMara | mMS | 1.144e-6 | 6.493e-6 |

### Supplementary Figures

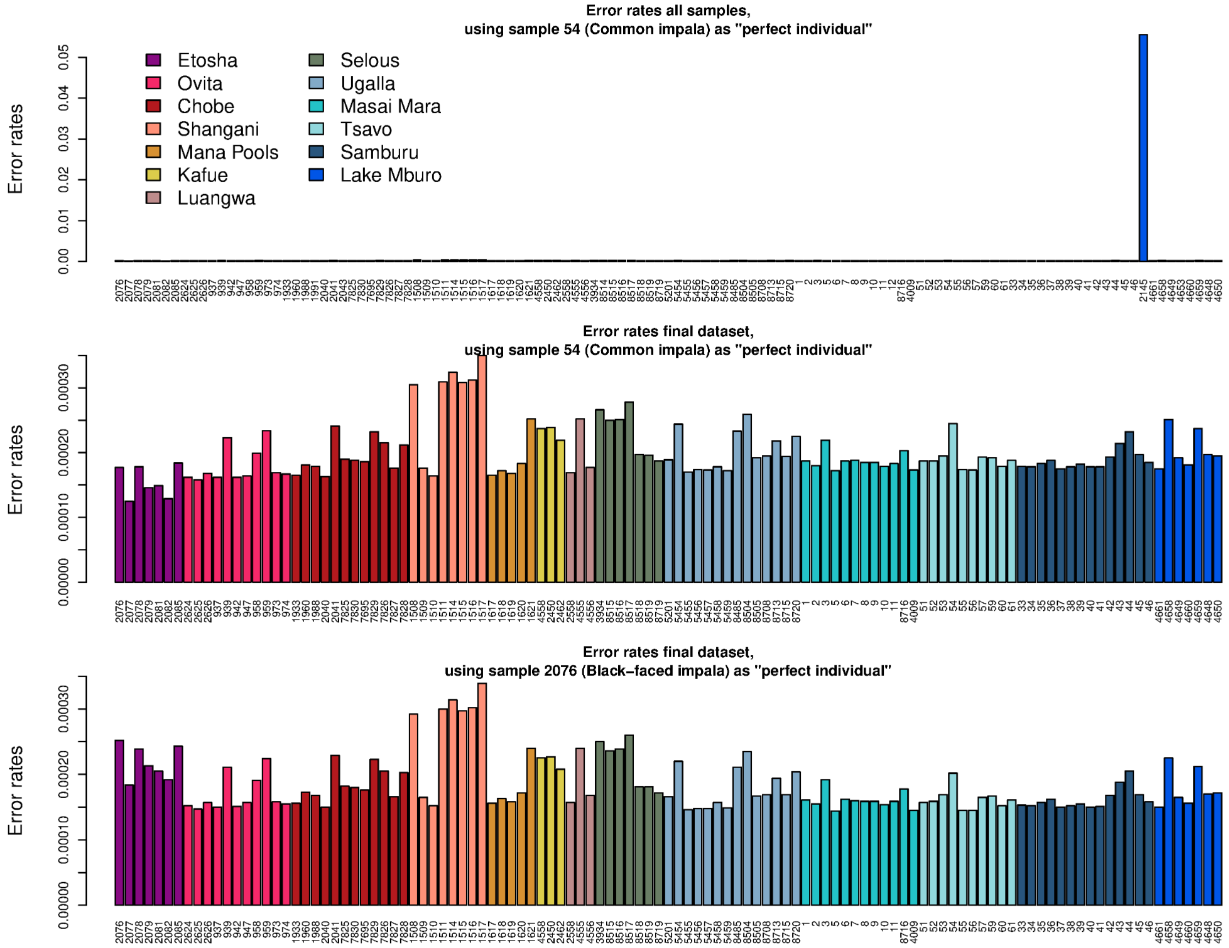

Supplementary Figure S1. Error rates estimated using the "perfect individual" method. The top two panels show error rate estimates when using a high depth common impala from Tsavo, Kenya, as the perfect individual with and without sample 2145, respectively. The bottom panel shows error rates using a high depth black-faced impala from Etosha, Namibia, as a perfect individual instead.

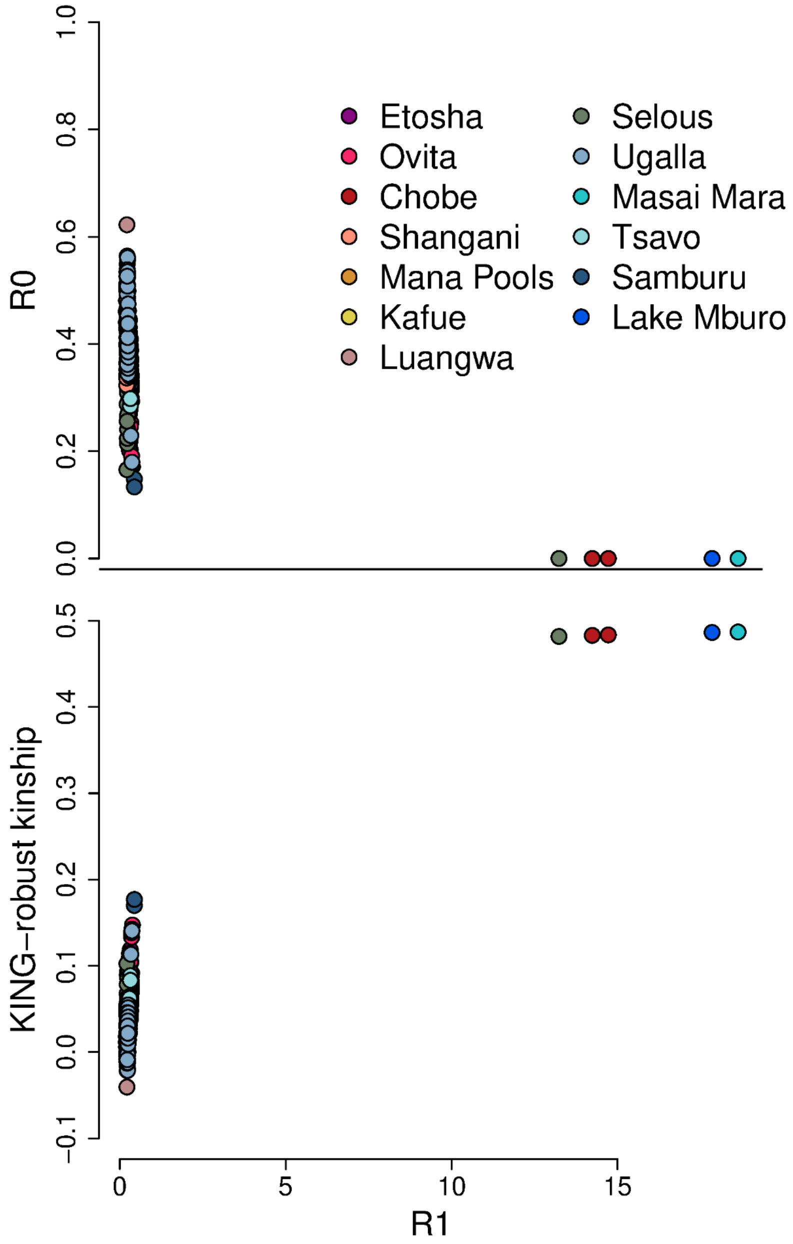

Supplementary Figure S2: Duplicate identification using an allele frequency free method (Waples et al. 2019). The plot shows, for each pair of samples from the same locality, the R0, R1 and KING-robust kinship, which are all based on ratios of identity by state sharing between pairs of individuals. While the estimates are too noisy to distinguish related from unrelated individuals, they allow clearly identifying 5 pairs of duplicate samples with extreme R1 values (>10), KING-robust of around 0.5 and R0 of 0.

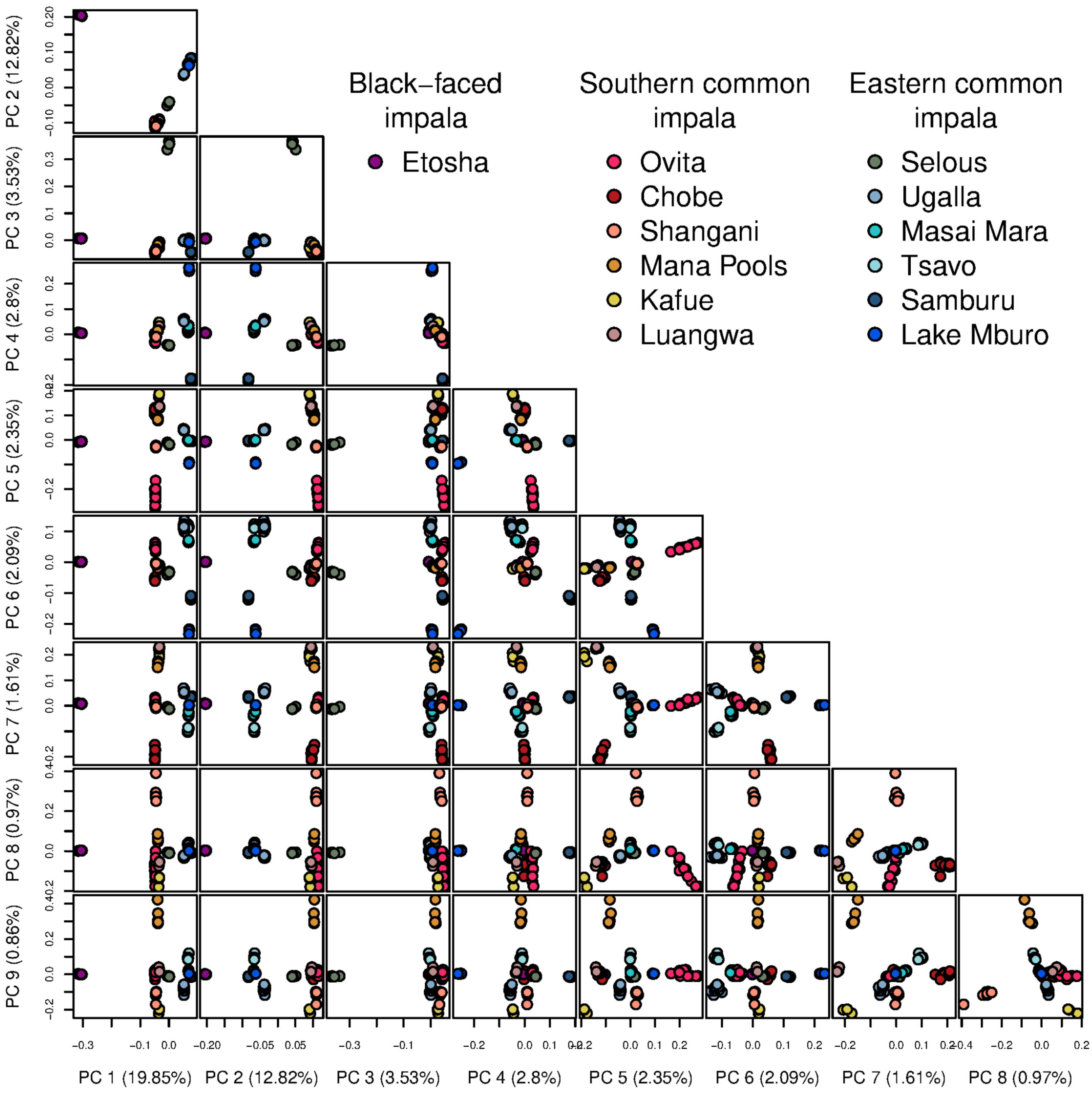

Supplementary Figure S3: PCA plots for all pairs of principal components from 1 to 9. The covariance matrix was estimated from genotype likelihoods with PCAngsd. The number in parenthesis on each axis is the percentage of the total variance explained by the corresponding PC.

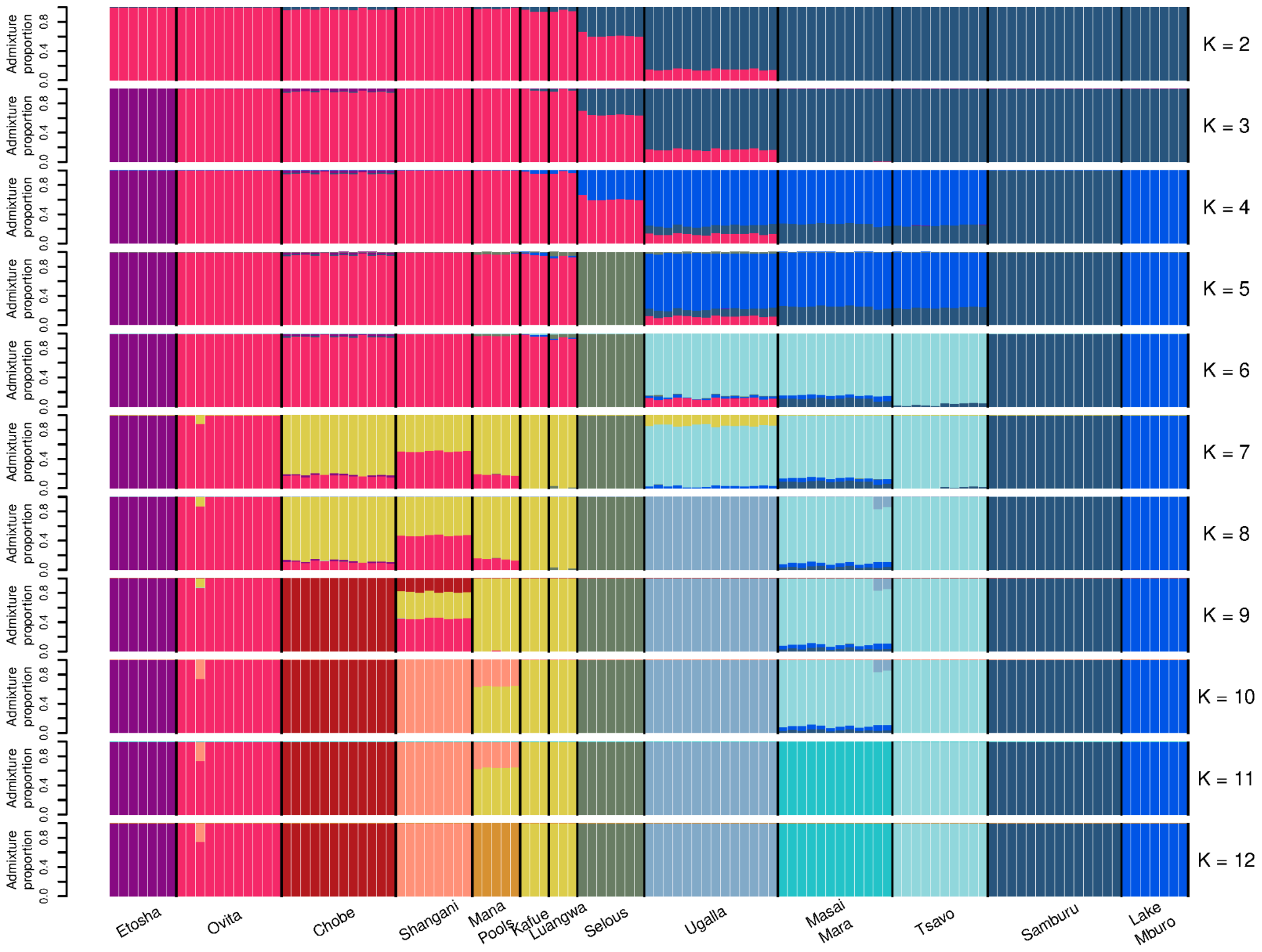

Supplementary Figure S4: Admixture proportions assuming the number of ancestral populations is K=2 to K=12 estimated from genotype likelihoods with NGSadmix.

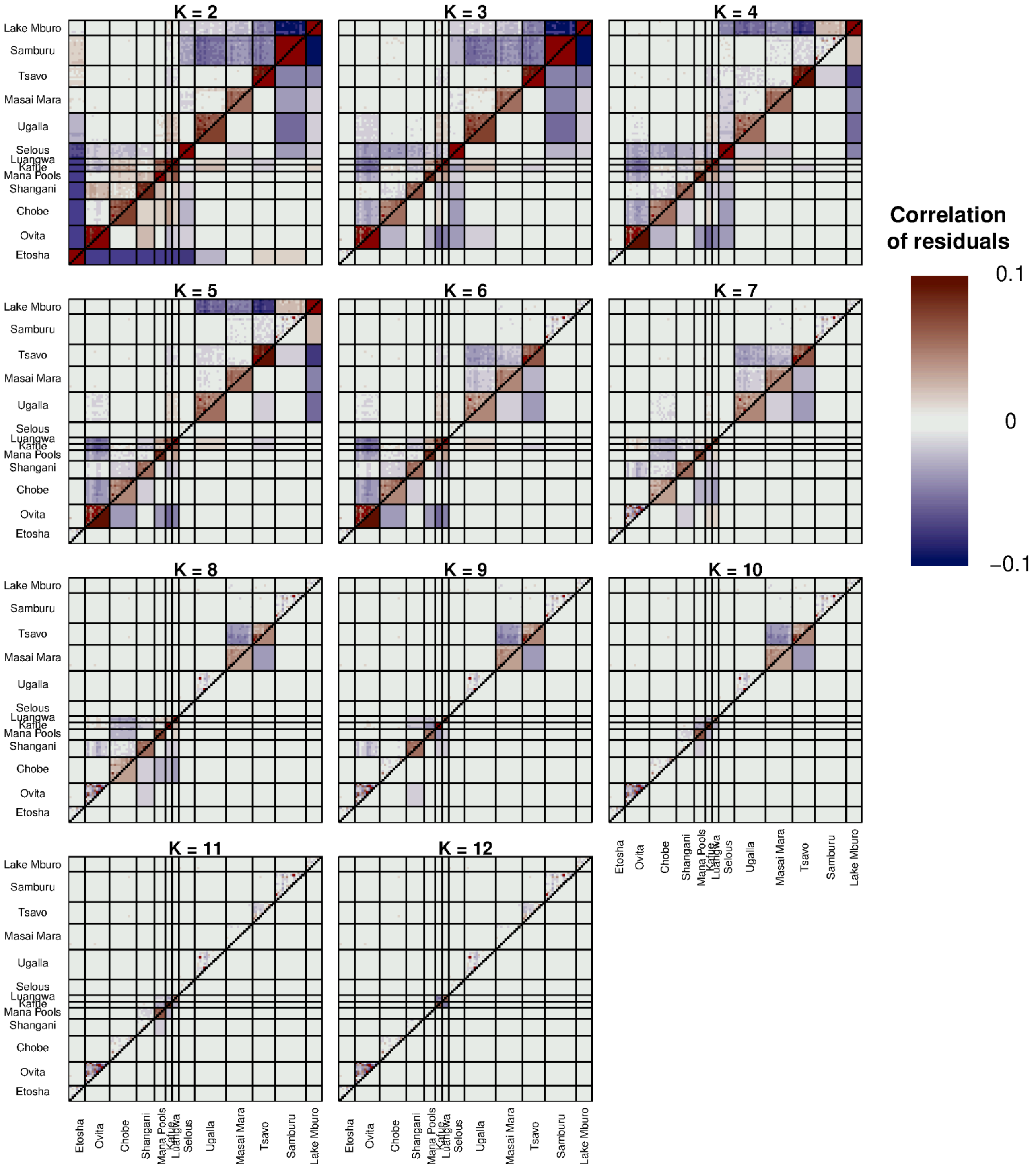

Supplementary Figure S5: Evaluation of the estimated admixture proportions (shown in Supplementary Figure S4) using the individual pairwise correlation of residuals (above diagonal), and population mean pairwise correlation of residuals (below diagonal), estimated with evalAdmix.

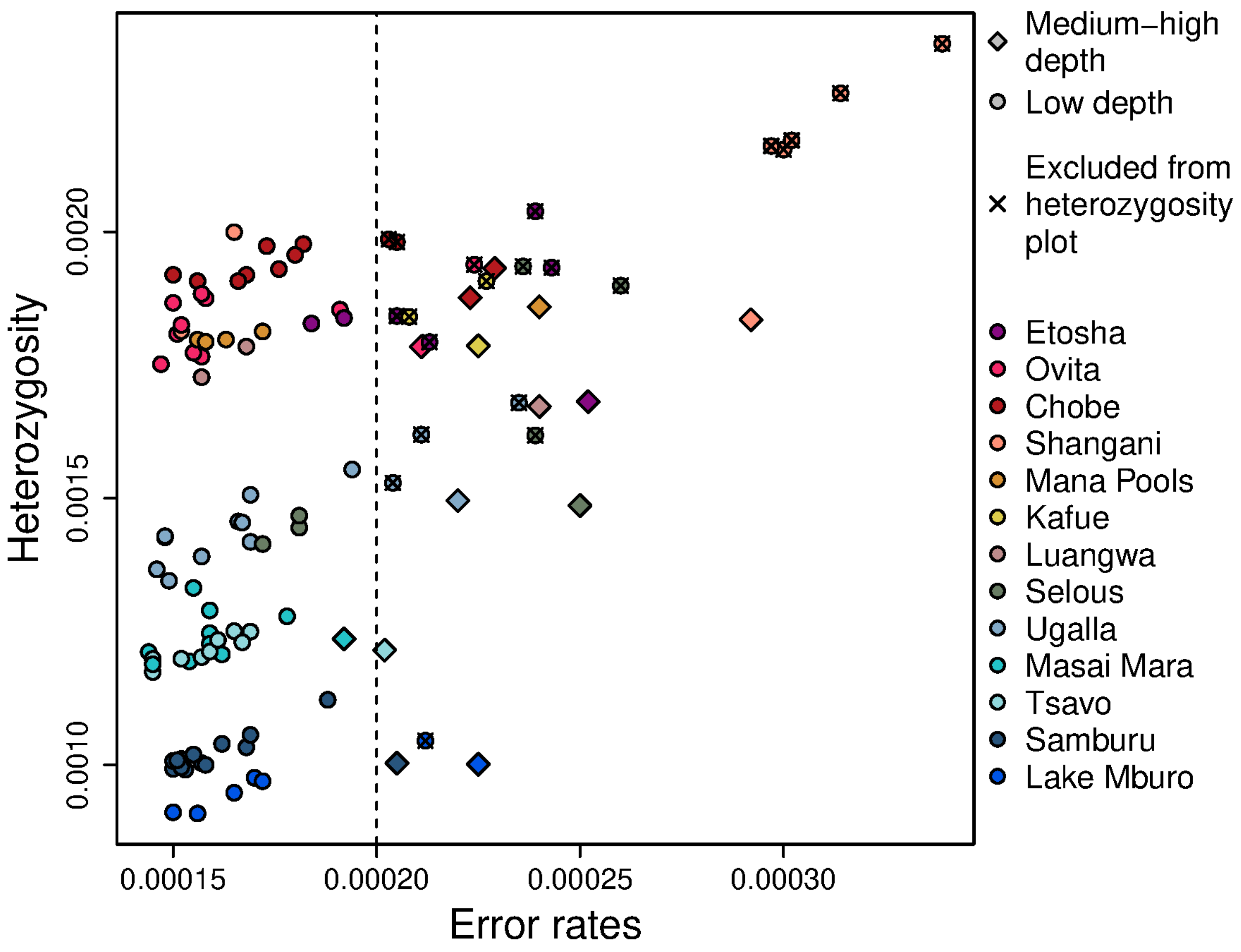

Supplementary Figure S6: Per-sample heterozygosity estimates versus the error rate estimated using sample 2076 as the “perfect individual”. Point colour indicates the sample’s locality of origin, while shape distinguishes between samples sequenced at low or high depth. Low-depth samples with high error rates showed elevated heterozygosity relative to samples from the same locality but with low error rates. In contrast, medium-high depth samples did not show elevated heterozygosity, even when their error rates were slightly higher. Based on this, low-depth samples with estimated error rates above 0.0002 are excluded from the heterozygosity boxplot in Figure 2A.

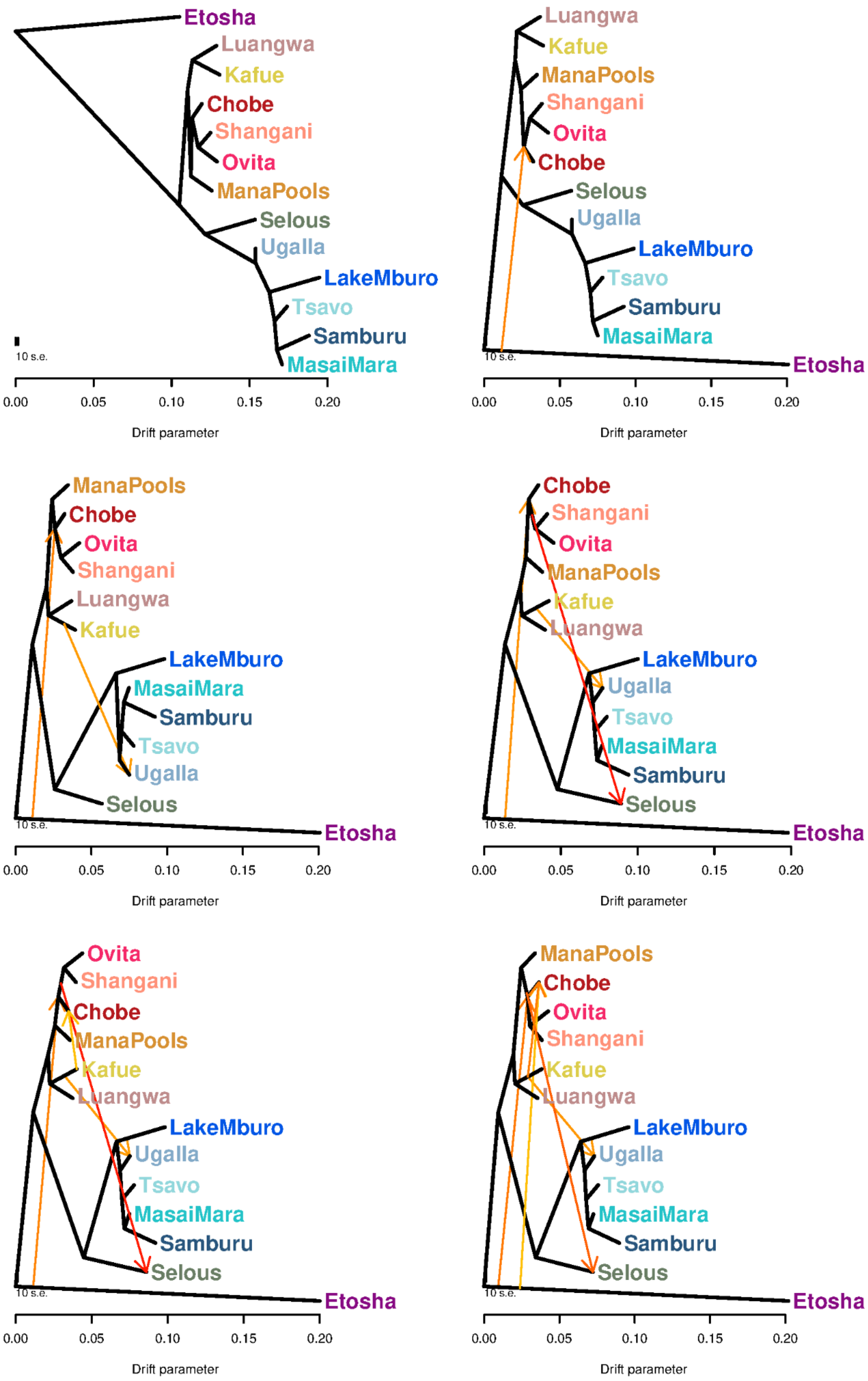

Supplementary Figure S7: Population trees inferred by TreeMix assuming 0 to 5 migration events.

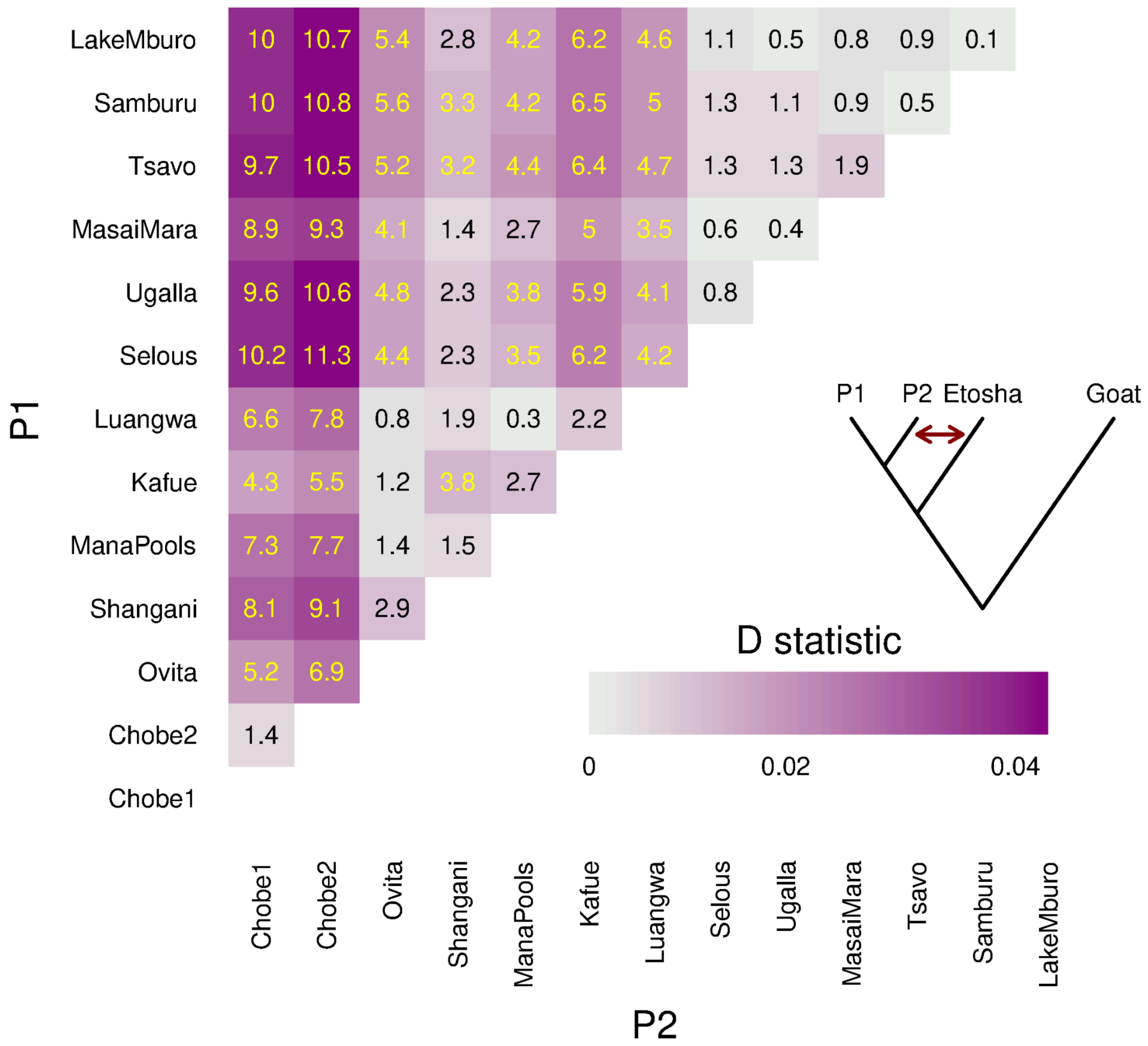

Supplementary Figure S8: Heatmap of D-statistics with the topology (((P1,P2), black-faced impala),Goat), where all possible pairs of the common impala populations are used as P1 and P2 and where black-faced impala is represented by the Etosha population.

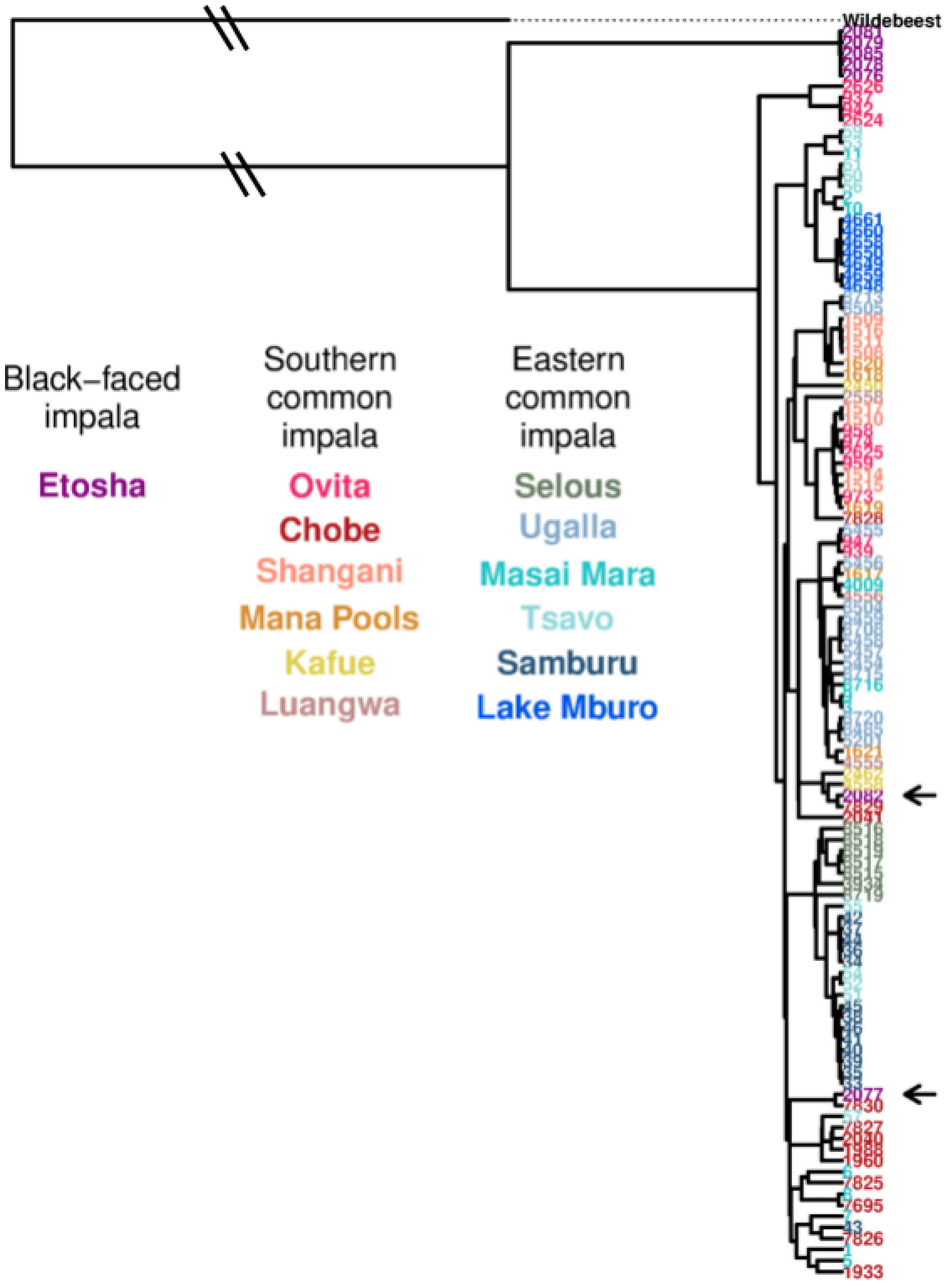

Supplementary Figure S9: mtDNA tree for all the impala samples. The branch length leading to the wildebeest is not scaled. Arrows indicate the two black-faced impalas from Etosha (Namibia) whose mtDNA clusters with common impalas from Chobe (Botswana).

Supplementary Figure S10: admixture graphs that fit equally well to the reduced dataset shown in Fig 3A. Attached as a separate file.

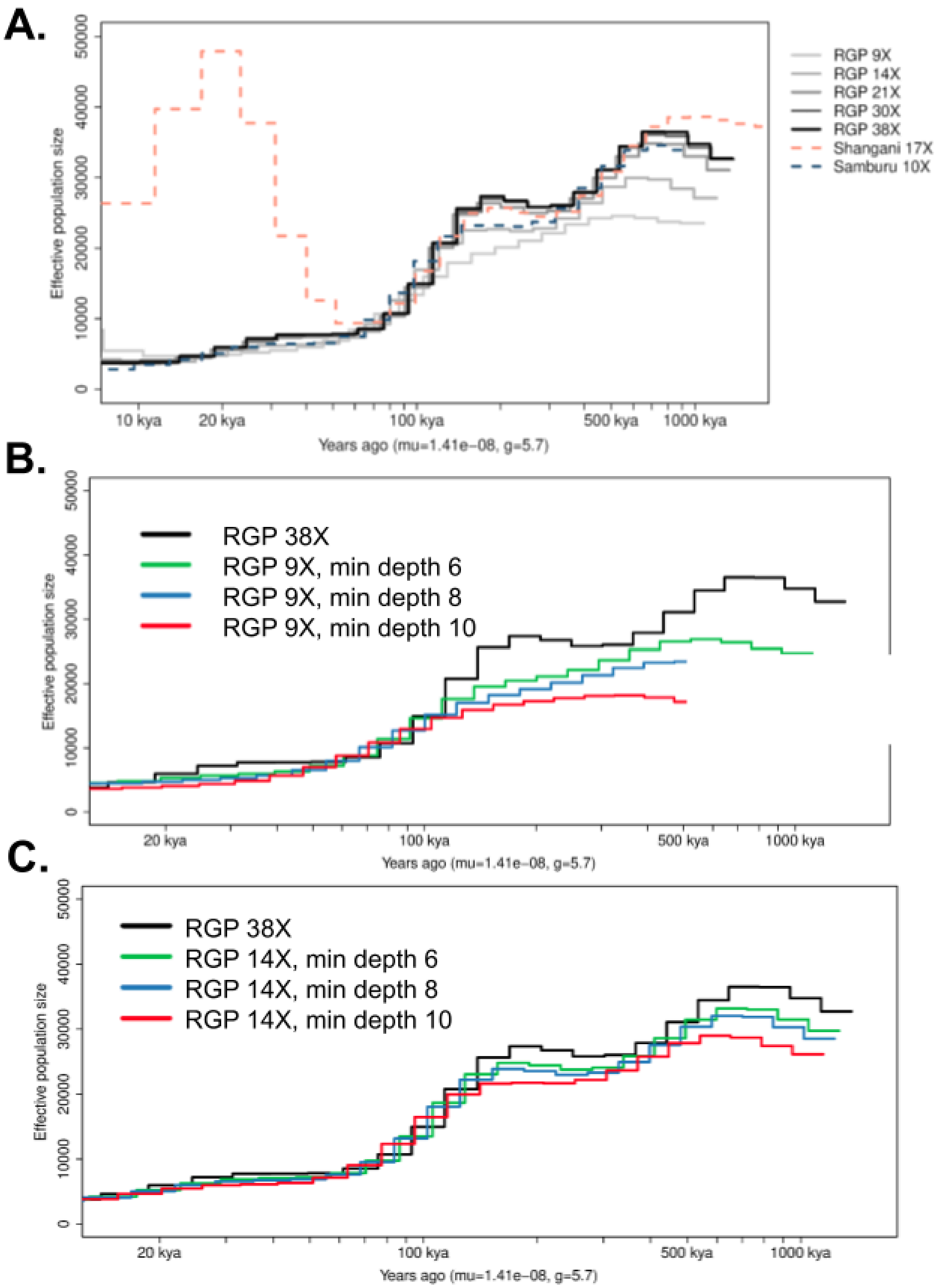

Supplementary Figure S11: Effect of sequencing depth and filtering on PSMC estimates. A. Effective population size trajectories over time inferred for the sample from the ruminant genome project (RGP), downsampled at different depths as well as for the high depth sample from Shangani and for the medium-depth sample from Samburu. This suggests that the differences in ancient time seen when plotting all samples in Fig. 4 are artefacts due to low sequencing depth. B. and C. Effect of different minimum depth thresholds on site inclusion for PSMC, when using the RGP sample subsampled to an average coverage of 9X (B.) and 14X (C.), with the RGP sample at 38X as a reference for the non depth biased Ne trajectory estimate. The depth bias is alleviated by using a less stringent minimum depth threshold, maximising thus the amount of sites included even though this would also increase the expected proportion of heterozygous sites lost.

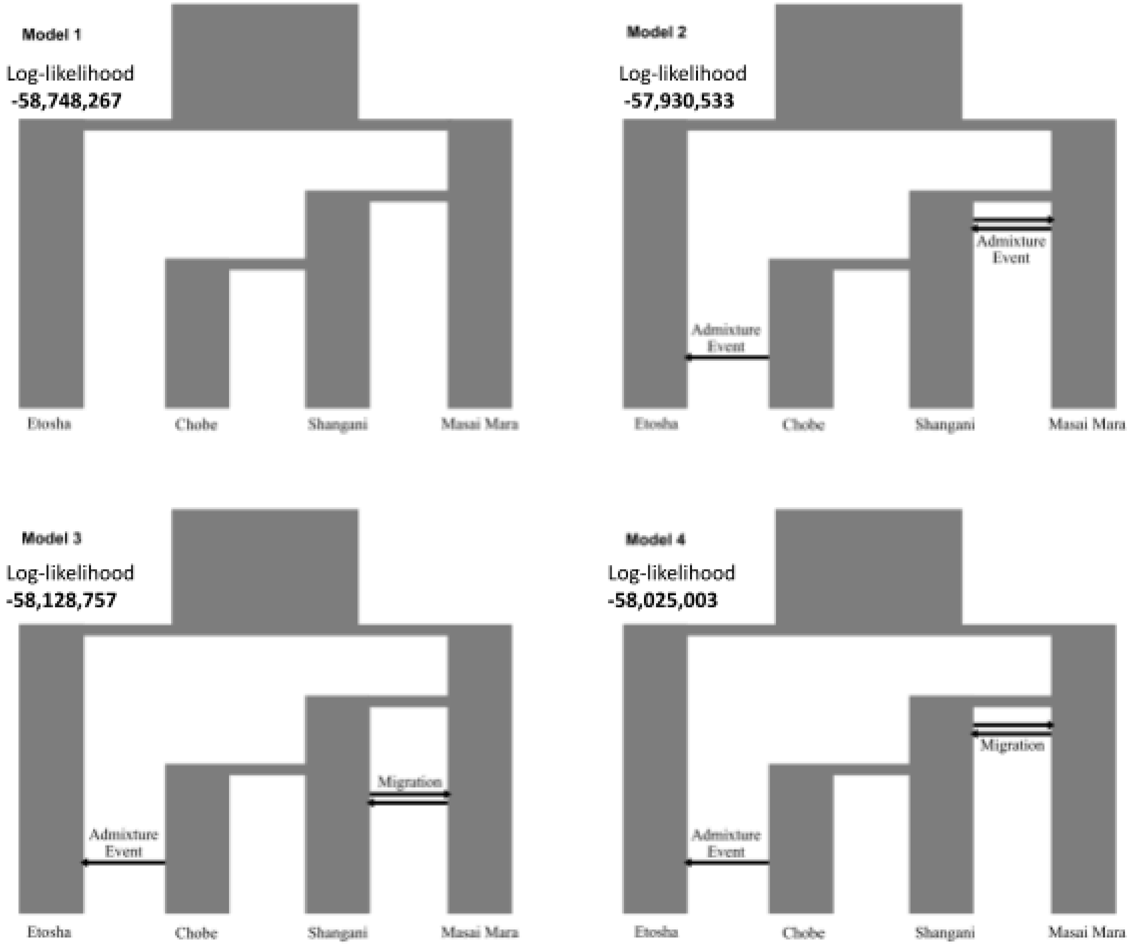

Supplementary Figure S12: Schematic visualisation of the 4 different demographic models tested with fastsimcoal2, together with the log likelihood of the maximum likelihood estimated parameters.

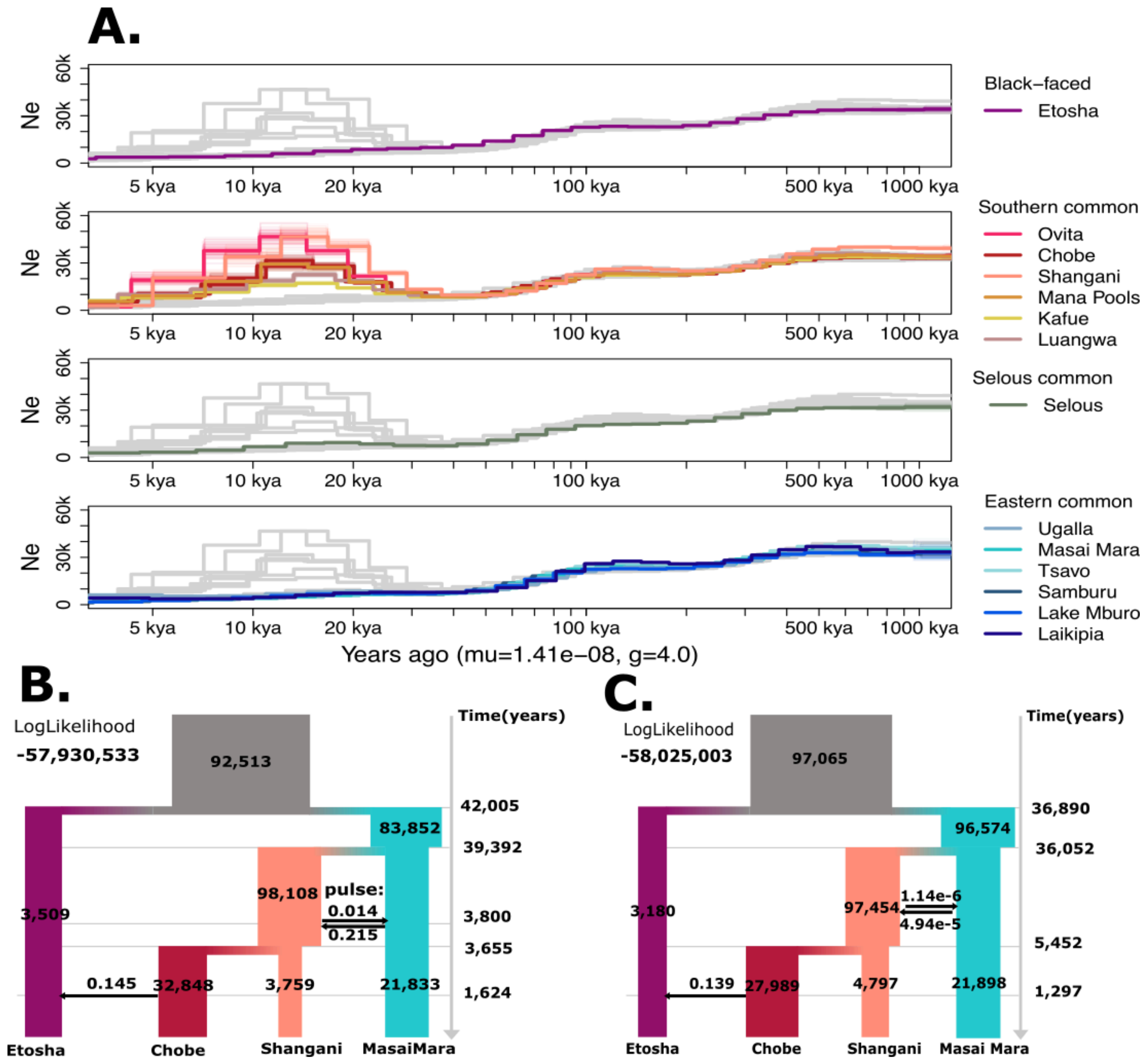

Supplementary Figure S13: A. Long-term effective population sizes for all impala populations, estimated with PSMC and B. and C. demographic history estimated with fastsimcoal2 with two models. Times are scaled from generations to years assuming a generation time of 4.03 (in contrast with generation time of 5.7 in Figure 4; see Methods).

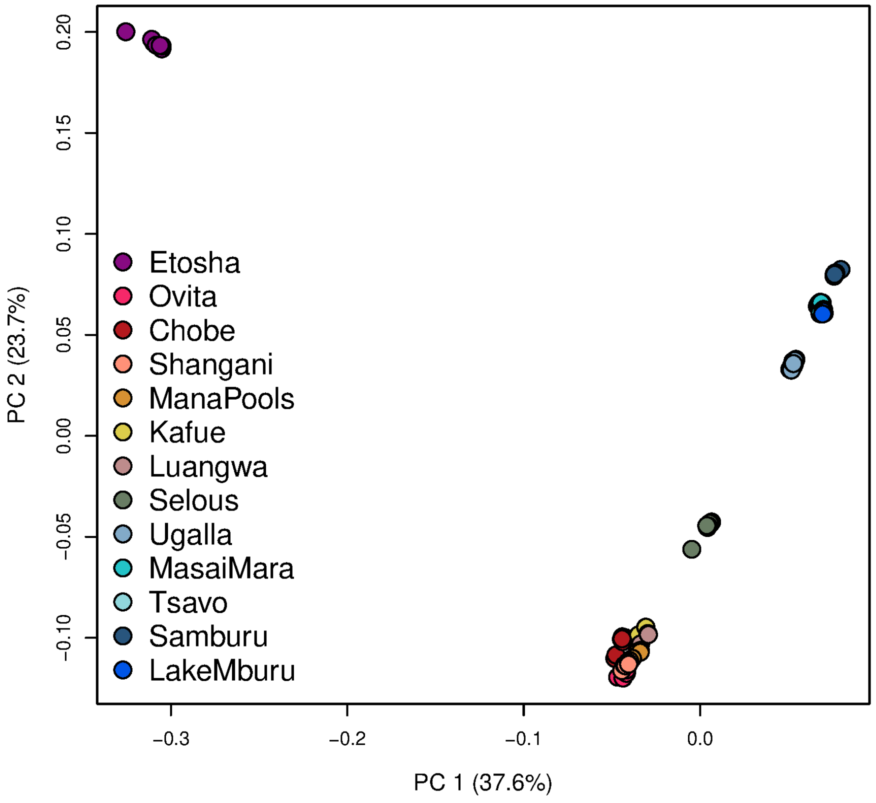

Supplementary Figure S14. PCA performed on the imputed dataset, showing almost identical results to the PCA from the genotype likelihoods in Figure 1B.

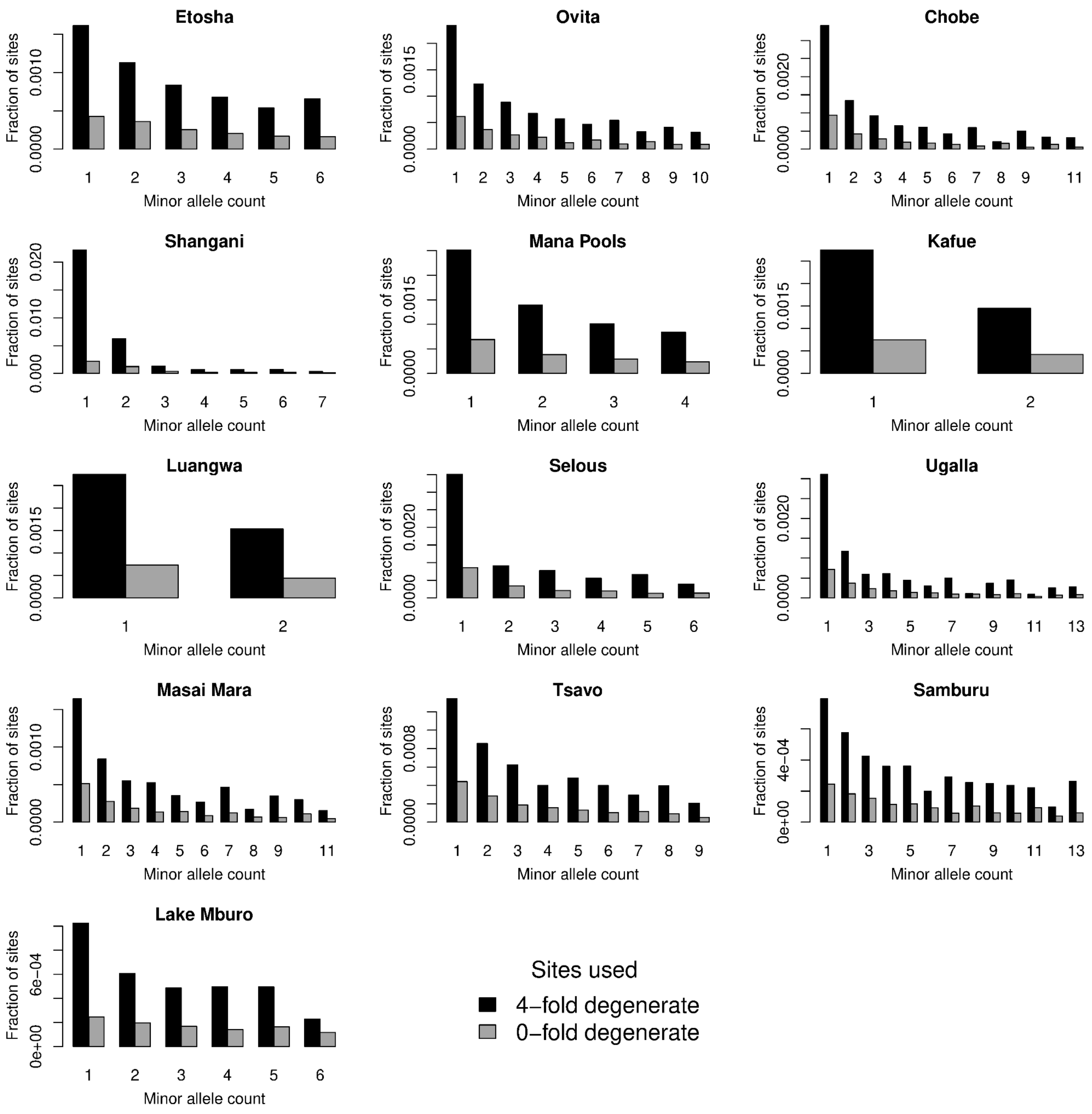

Supplementary Figure S15: Estimated SFS in 0 fold and in 4 fold degenerate sites for all impala populations. More than 99% of the sites for all the populations are in the fixed category bin (minor allele count = 0) which is not shown on the plot.

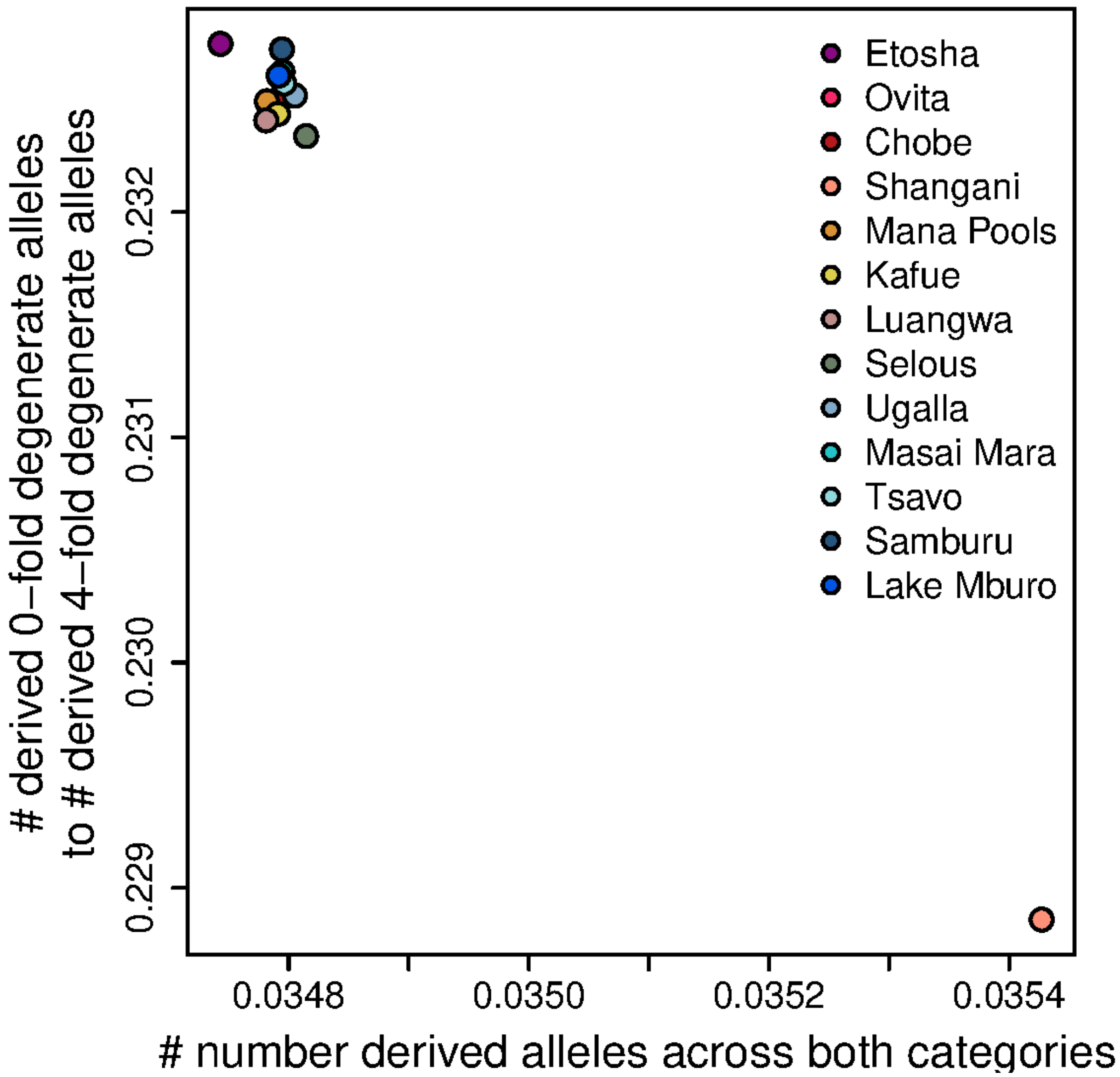

Supplementary Figure S16: Effect of excess error rates on the efficiency of selection analyses. Total number of derived alleles plotted against the ratio of derived alleles in 0-fold to 4-fold degenerate sites. The Shangani population is excluded from the boxplot in Fig. 5 based on this.

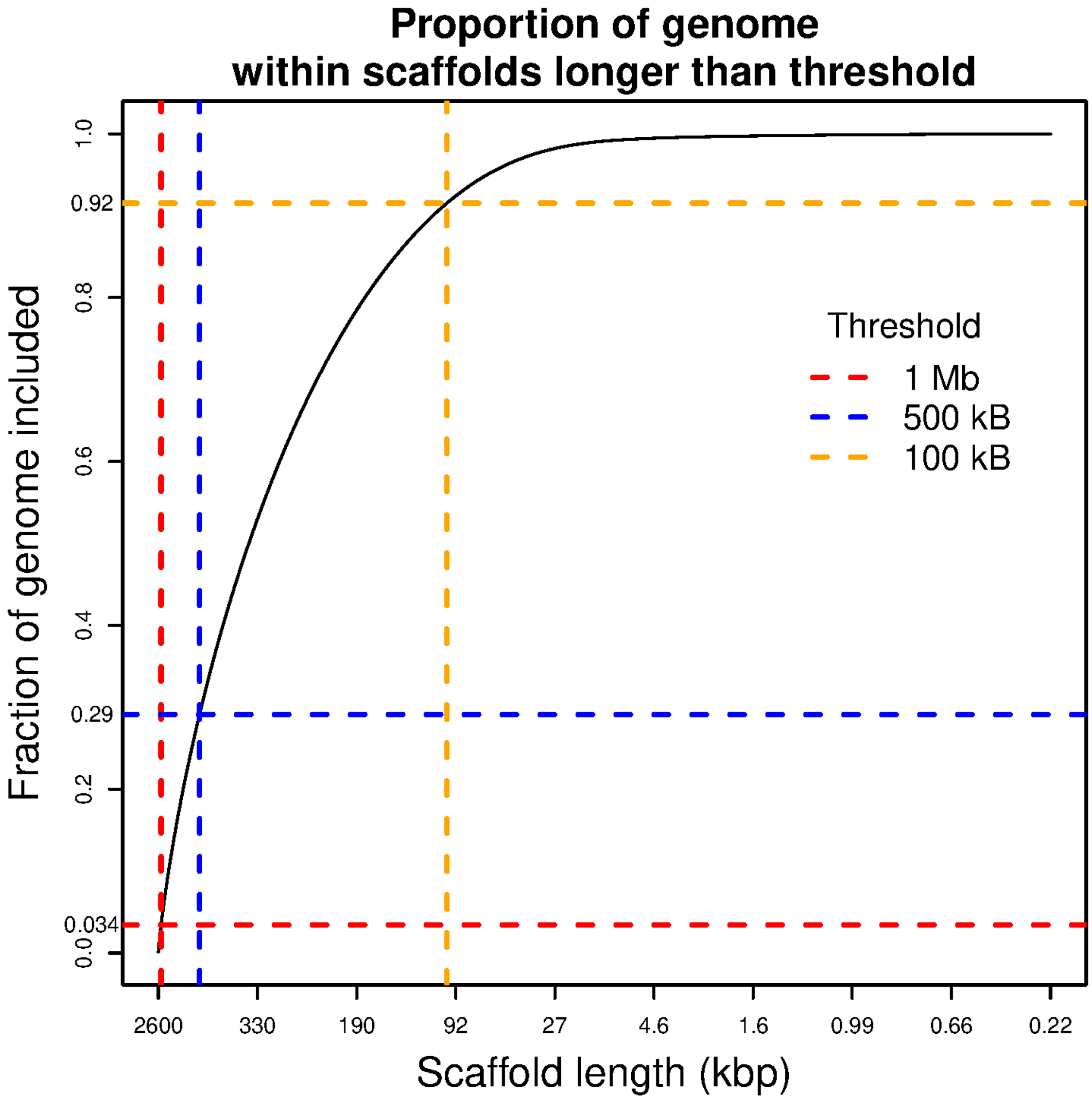

Supplementary Figure S17: Fraction of the impala reference genome retained as a function of the minimum scaffold length required to not filter out scaffolds. Based on the plot a minimum scaffold length of 100 kb was set.
