## supplementary figure s10 for "Extensive population structure highlights an apparent paradox of stasis in the impala (*Aepyceros melampus*)"

idx: 25 Best model. adm: 1 Scoretest: 64.4223303248011

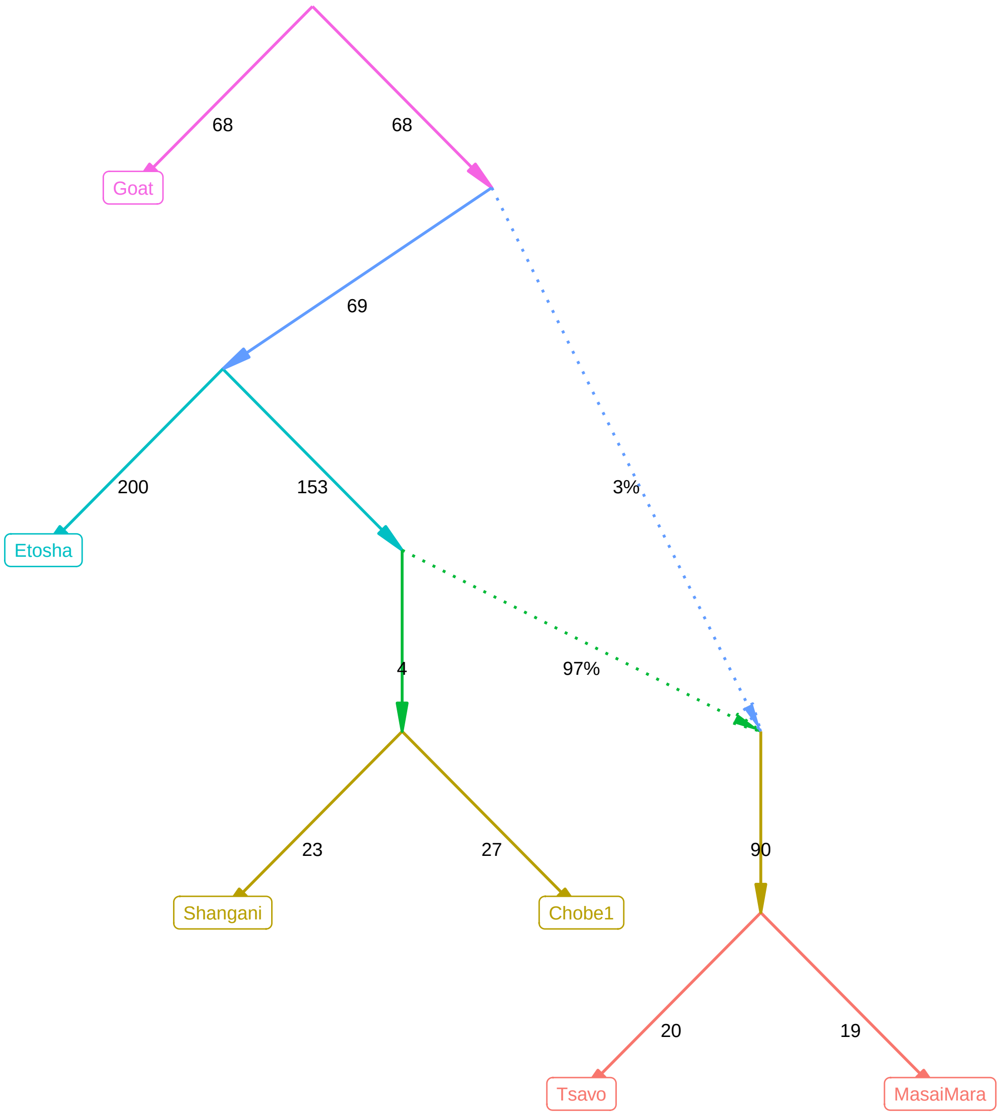

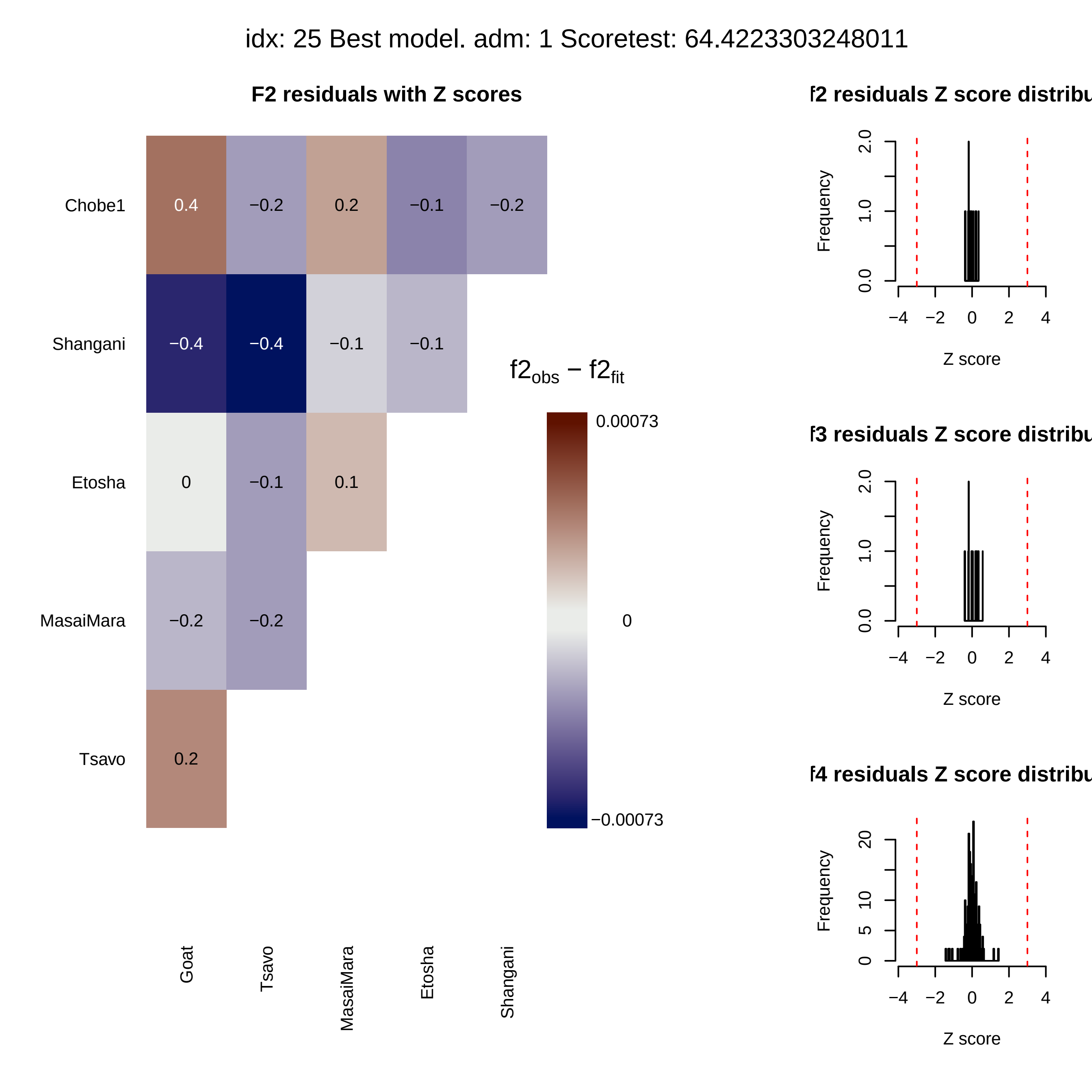
idx: 25 Best model. adm: 1 Scoretest: 64.4223303248011
F2 residuals with Z scores
Chobe1
Shangani
Etosha
MasaiMara
Tsavo
0.4
−0.2
0.2
−0.1
−0.2
−0.4
−0.4
−0.1
−0.1
0
−0.1
0.1
−0.2
−0.2
0.2
Goat
Tsavo
MasaiMara
Etosha
Shangani
f2obs − f2fit
0.00073
0
−0.00073
f2 residuals Z score distribu
f3 residuals Z score distribu
f4 residuals Z score distribu
Frequency
Z score

idx: 24 pval to best: 0.7568 adm: 1 Score_test2: 65.4874008861579 diff: −1.0651

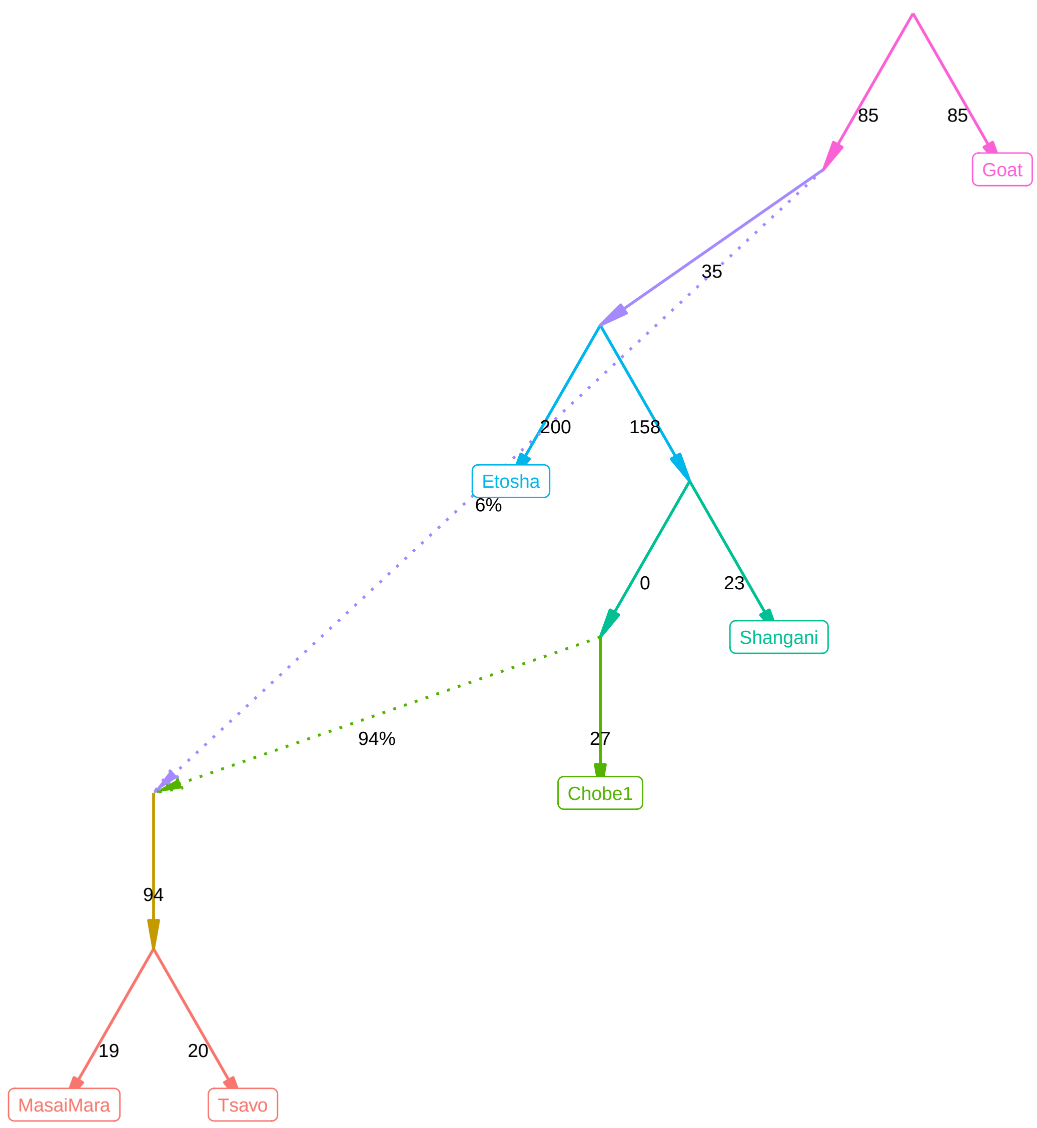

idx: 24 pval to best: 0.7568 adm: 1 Score_test2: 65.4874008861579 diff: −1.0651

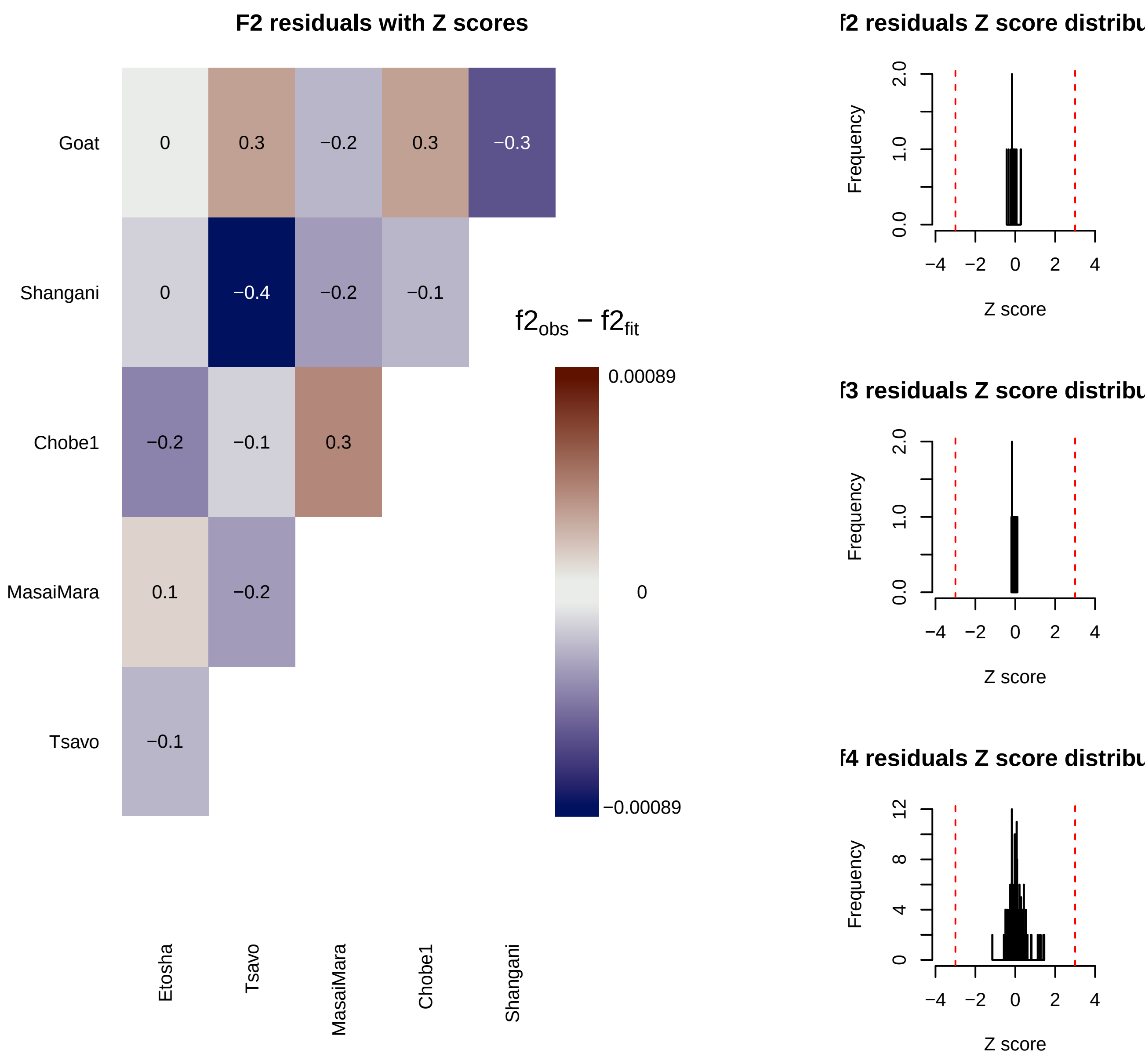

idx: 23 pval to best: 0.8447 adm: 1 Score_test2: 65.5606333995951 diff: −1.1383

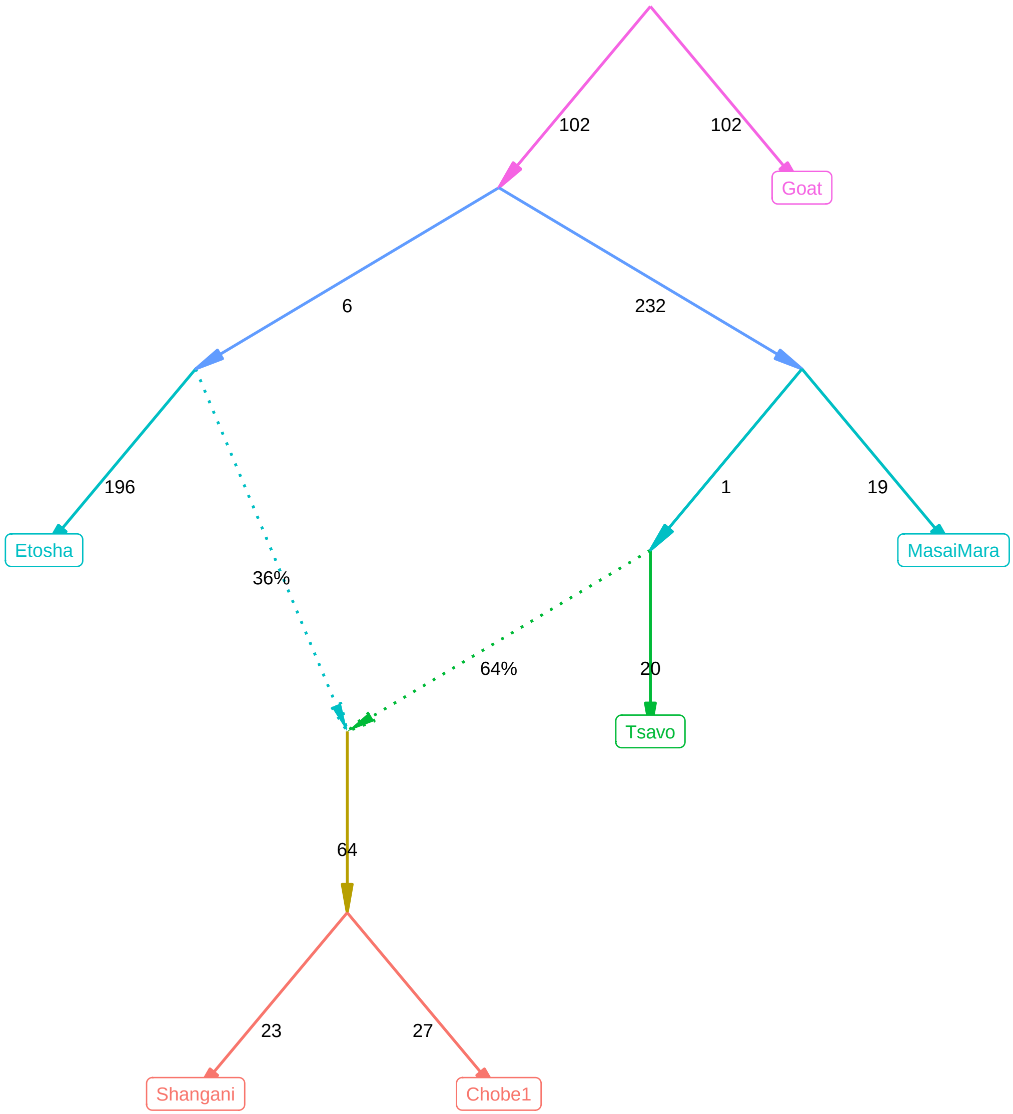

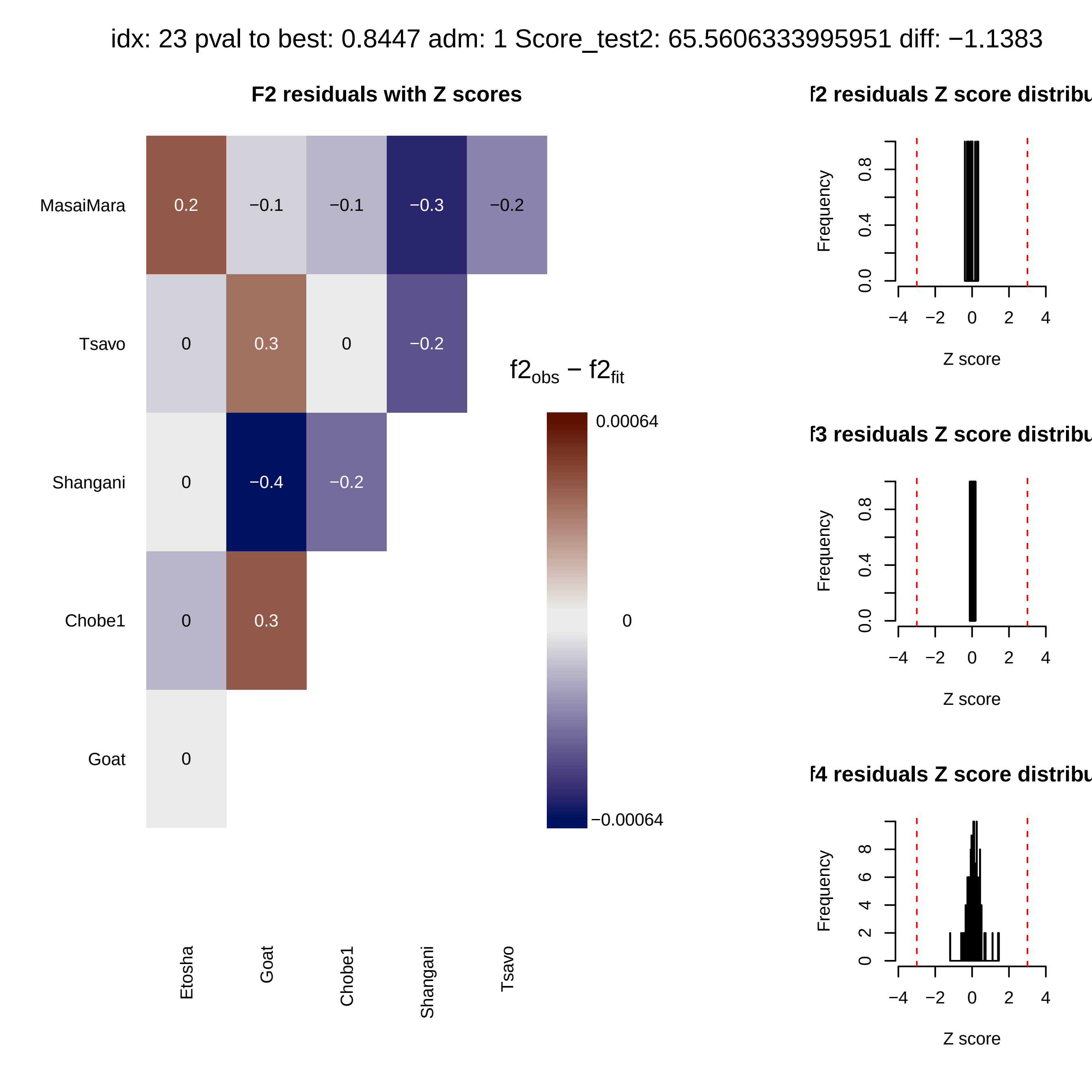

idx: 23 pval to best: 0.8447 adm: 1 Score_test2: 65.5606333995951 diff: −1.1383
F2 residuals with Z scores
MasaiMara
Tsavo
Shangani
Chobe1
Goat
0.2
−0.1
−0.1
−0.3
−0.2
0
0.3
0
−0.2
0
−0.4
−0.2
0
0.3
0
Etosha
Goat
Chobe1
Shangani
Tsavo
f2obs − f2fit
0.00064
0
−0.00064
f2 residuals Z score distribu
f3 residuals Z score distribu
f4 residuals Z score distribu
Frequency
Z score

idx: 22 pval to best: 0.8032 adm: 1 Score_test2: 65.83380344284 diff: −1.4115

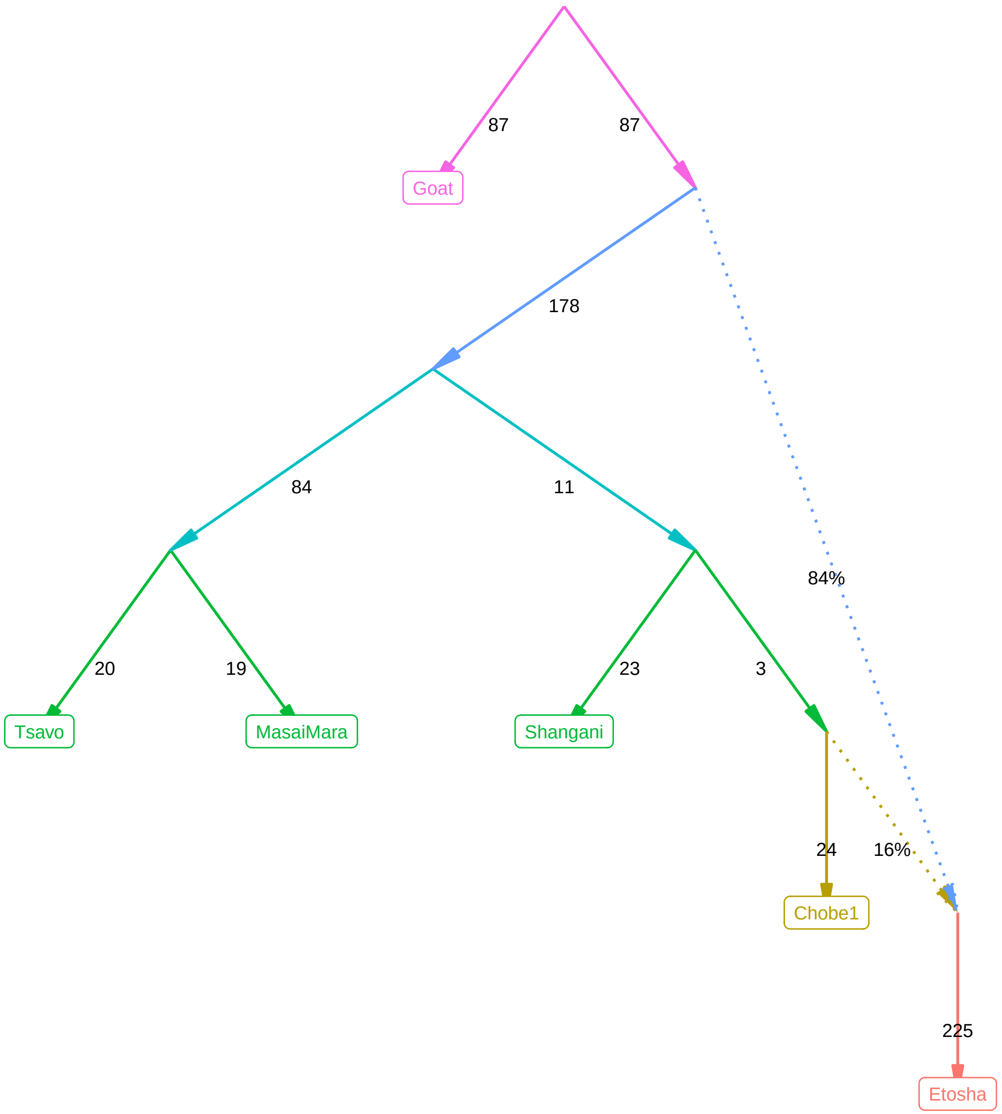

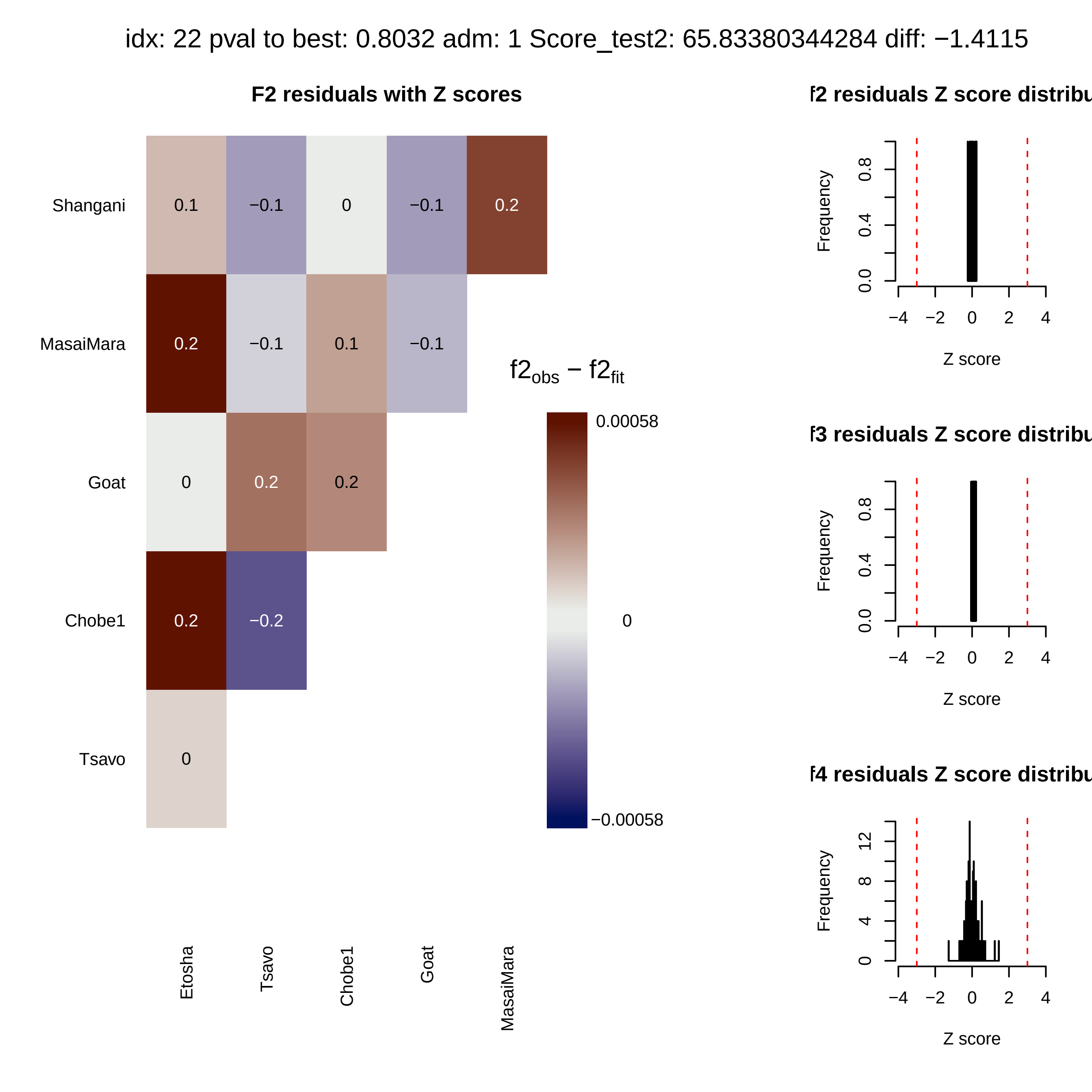

idx: 22 pval to best: 0.8032 adm: 1 Score_test2: 65.83380344284 diff: −1.4115
F2 residuals with Z scores
Shangani
MasaiMara
Goat
Chobe1
Tsavo
0.1
−0.1
0
−0.1
0.2
0.2
−0.1
0.1
−0.1
0
0.2
0.2
0.2
−0.2
0
Etosha
Tsavo
Chobe1
Goat
MasaiMara
f2obs − f2fit
0.00058
0
−0.00058
f2 residuals Z score distribu
f3 residuals Z score distribu
f4 residuals Z score distribu
Frequency
Z score
